# Spatial Control of Draper Receptor Signaling Initiates Apoptotic Cell Engulfment

**DOI:** 10.1101/198986

**Authors:** Adam P. Williamson, Ronald D. Vale

## Abstract

The engulfment of apoptotic cells is essential for tissue homeostasis and responding to damage. Engulfment is mediated by receptors that recognize ligands exposed on apoptotic cells, such as phosphatidylserine (PS). Here, we convert Drosophila S2 cells into proficient phagocytes by transfecting the Draper engulfment receptor and replacing apoptotic cells with PS-coated beads. We show that PS-ligated Draper forms microclusters that exclude a bulky transmembrane phosphatase and recruit phosphotyrosine binding proteins, revealing a triggering mechanism similar to the T cell receptor (TCR). Analogous to the TCR, Draper’s extracellular domain and PS can be replaced with FRB and FKBP respectively, resulting in a rapamycin-inducible engulfment system. However, in contrast to the TCR, we show that localized signaling at Draper microclusters results in time-dependent depolymerization of actin filaments. Collectively, our results reveal mechanistic similarities and differences between the receptors involved in apoptotic corpse clearance and mammalian immunity and demonstrate that engulfment can be reprogrammed towards non-native targets.

**Condensed title:** Spatial Control of Engulfment Signaling

## Introduction

The prompt clearance of dying cells and debris is essential for maintaining homeostasis and promoting tissue repair (Reddien and Horvitz, 2004; Neumann et al., 2015; Arandjelovic and Ravichandran, 2015). In healthy tissue, resident and infiltrating phagocytes clear cell corpses and debris through specific recognition, uptake, and digestion (Elliott and Ravichandran, 2010). Defects in clearance result in autoimmunity and further tissue damage (Elliott and Ravichandran, 2010; Iram et al., 2016; Kawano and Nagata, 2018). Despite the importance of efficient clearance across multicellular life, mechanisms of engulfment receptor activation remain poorly understood in comparison to other signaling systems. Defining the molecular basis of engulfment receptor activation could lead to new strategies for enhancing clearance under conditions of extreme injury or programming phagocytes to eliminate functionally-relevant targets such as cancer cells or pathogens.

The initial event in apoptotic cell clearance involves interactions of “eat me” ligands exposed on dying cells with receptors on the phagocyte. Phosphatidylserine (PS) exposed on the outer leaflet of the plasma membrane constitutes one such “eat me” ligand, although several other protein ligands likely participate as well (Fadok et al., 1992; Segawa and Nagata, 2015; Ravichandran and Lorenz, 2007). Ligand binding causes receptor phosphorylation, a process known as “receptor triggering”, and this event leads to cytosolic signaling that ultimately promotes cytoskeletal rearrangements that power corpse internalization (Ravichandran and Lorenz, 2007; Reddien and Horvitz, 2004). For most transmembrane receptors, triggering proceeds via one of two mechanisms: ligand induced receptor conformational change to transmit the signal across plasma membranes or “kinetic segregation”, in which spatially organized zones of ligated receptors physically exclude phosphatases to favor net receptor phosphorylation and activation. Epidermal Growth Factor Receptor (EGFR) and G Protein-Coupled Receptors (GPCRs) are examples of conformation-induced activation (Dawson et al., 2005; Erlandson et al., 2018), while the mammalian immune receptors that promote T cell activation and Fc receptor-dependent engulfment of opsonized targets are triggered via a kinetic segregation mechanism (Davis and van der Merwe, 2006; Freeman et al., 2016). It remains unclear which activation mechanism corpse clearance receptors utilize to transmit the “eat me” signal across phagocyte plasma membranes. Here, we use receptor triggering as an inlet to address two open questions about engulfment signaling initiation: Do apoptotic ligands transmit a signal across the plasma membrane via a receptor conformational change or a kinetic segregation mechanism? How are ligated receptors organized on phagocyte plasma membranes to potentiate engulfment signaling?

To gain insight into these questions, we focused on Draper, a Drosophila engulfment receptor. Draper is expressed in glia, where it promotes clearance of damaged axons, and in the somatic epithelium, where it functions to remove dying cells from the follicle (Freeman et al., 2003; MacDonald et al., 2006; Etchegaray et al., 2012). Draper is similar in domain structure to the mammalian protein Megf10, and to CED-1, the first described apoptotic corpse receptor in C. elegans. All three receptors contain single pass transmembrane domains that connect extracellular EGF repeats to intracellular tyrosine phosphorylation sites that are presumed to recruit one or more downstream effectors (Scheib et al., 2012; Ziegenfuss et al., 2008; Zhou et al., 2001). Draper function has been dissected genetically, especially in its damaged axon clearance role, which defined a “parts list” of other proteins involved in engulfment signaling, including the kinases Src42a and Shark and the adapter protein Crk (Ziegenfuss et al., 2012; 2008; Lu et al., 2014). However the mechanism by which receptor ligation leads to effector recruitment remains poorly understood.

Here, we have reconstituted Draper-dependent engulfment in Drosophila S2 cells to dissect receptor triggering. We find that the lipid phosphatidylserine (PS) incorporated into supported lipid bilayers on beads is sufficient to induce receptor phosphorylation and a signaling cascade leading to engulfment. This system allows a dramatic reduction in the complexity of the apoptotic cell as the engulfment target. Similar to the T Cell Receptor (TCR), ligated Draper forms mobile microclusters that shift a kinase-phosphatase balance toward receptor phosphorylation. However, unlike TCR microclusters, Draper microclusters locally deplete filamentous-actin (F-actin). By mass spectrometry, we further demonstrate that phosphorylation of Draper is ordered and full activation of Draper requires an initial ITAM phosphorylation followed by the modification of other tyrosines in its tail domain. We also reveal that the extracellular module of Draper and PS can be replaced with an artificial receptor-ligand interaction. Our reconstitution of the initial steps of Draper signaling provides new insight into the mechanism of engulfment receptor triggering and a strategy to reprogram engulfment to new targets.

## Results

### Cellular reconstitution of Draper-dependent engulfment

We used cultured Drosophila S2 cells, which are derived from a hemocyte lineage (Schneider, 1972), as a cellular system for reconstituting corpse clearance. Drosophila S2 cells display a low level of engulfment of cell corpses (Fig. 1A). We hypothesized that the level of engulfment might be low because S2 cells lack sufficient levels of engulfment receptors. For example, Draper is expressed at low levels in S2 cells as measured by RNA-Seq (Cherbas et al., 2011). To test whether introducing an exogenous engulfment receptor was sufficient to promote engulfment, we transfected S2 cells with Draper and assayed internalization of cell corpses using microscopy. Draper transfection increased the uptake of apoptotic cell corpses by ∼5 fold (Fig. 1A).

**Figure 1:**
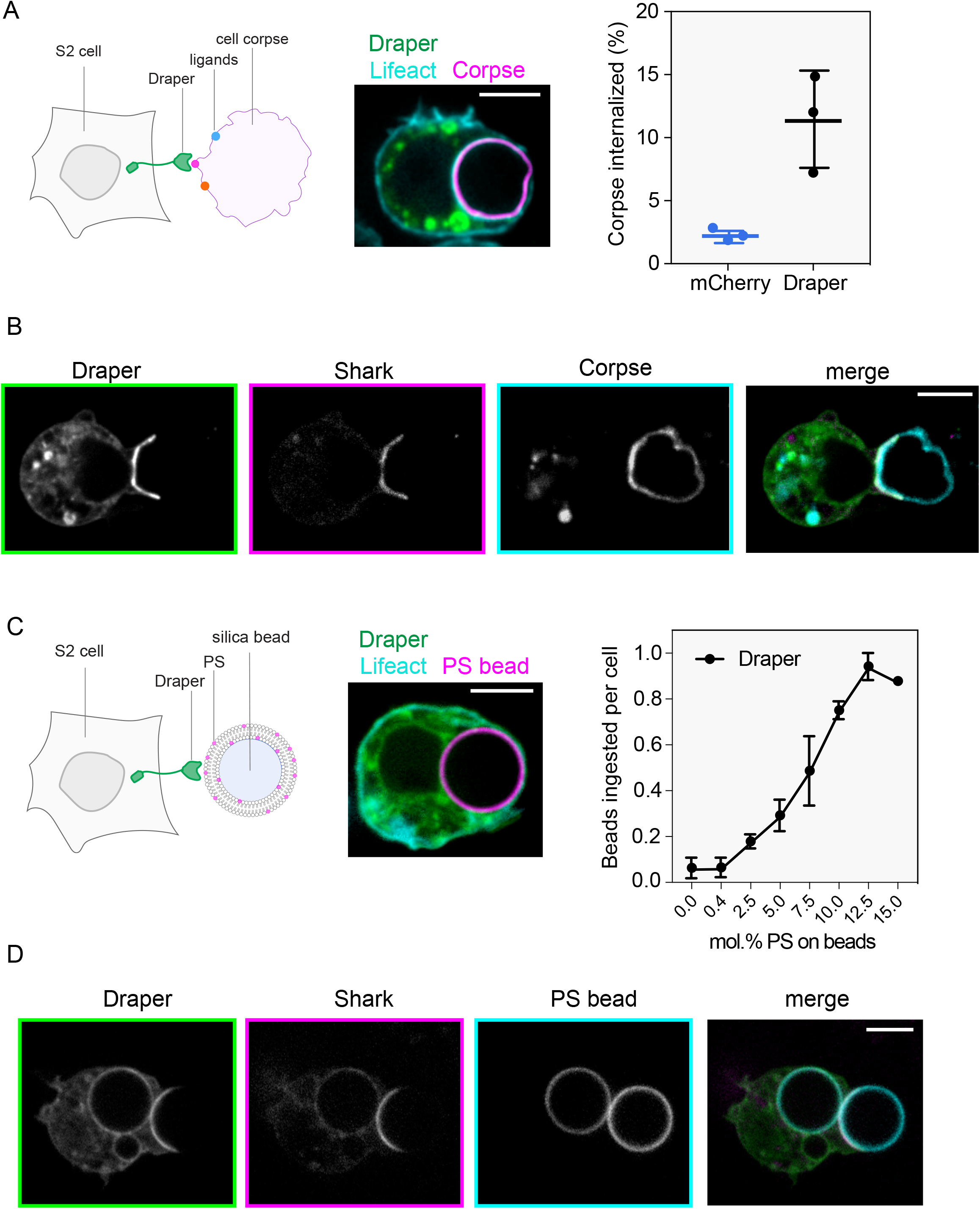
Cellular reconstitution of Draper-dependent engulfment. **A.** Schematic of apoptotic cell clearance by professional phagocytes; multiple receptors including Draper (green) recognize distinct ligands to promote engulfment of dying cell corpses (purple). Representative image of an Annexin V-APC labeled cell corpse fully internalized by a Draper-GFP expressing S2 cell. Overexpression of Draper-GFP in S2 cells results in a ∼5 fold increase in phagocytic proficiency toward apoptotic cell corpses. Draper-GFP+ S2 cells were incubated with labeled cell corpses and phagocytic proficiency was assessed after 45 min incubation at 27° C. Results from 3 independent biological replicates are shown (mean ± SD). **B.** Draper-GFP’s ability to bring Shark to the synapse between cell corpse (cyan) and S2 cell was assessed by live cell imaging. Draper-GFP signal (left) overlaps with Shark-mCherry localization (middle) at the synapse between corpse and S2 cell. Scale bar indicates 5 µm. **C.** Schematic of reconstitution system to study Draper signaling. The cell corpse was replaced with a silica bead coated with a supported lipid bilayer containing phosphatidylserine (PS). Representative image of an atto390-labeled 10% PS bead fully internalized by a Draper-GFP expressing S2 cell. Draper-GFP expressing S2 cells were incubated with beads containing increasing mol. % of PS in the supported lipid bilayer and phagocytic proficiency was assessed after 30 min incubation at 27° C. The mean number of beads fully ingested per cell for three independent biological replicates are shown (mean ± SD). **D.** Shark kinase is enriched at the synapse between 10% PS bead and the S2 cell. S2 cells were co-transfected with Draper-GFP and Shark-mCherry and incubated with 10% PS beads labeled with 0.5% atto390-DOPE. After 15 min, cells with active synapses were imaged by spinning disk confocal microscopy. The middle slice from a z-stack is shown. Scale bar indicates 5 µm.

The synapse between Draper transfected S2 cells and corpses also served as a site of local receptor activation. After engaging with a damaged axon, Draper expressed in glial cells becomes tyrosine phosphorylated at an ITAM motif, which then enables it to recruit a Syk-related tyrosine kinase called Shark (Ziegenfuss et al., 2008). Consistent with the studies of Ziegenfuss et al. (2008), we find a specific recruitment of Shark-mCherry to the plasma membrane to the interface between the corpse and S2 cell (Fig. 1B). Therefore ligation of Draper to one or more ligands on a cell corpse is sufficient to trigger the local recruitment of the cytosolic tandem SH2-domain containing kinase Shark and induce engulfment.

Apoptotic cells express an array of “eat me” ligands (Ravichandran and Lorenz, 2007). As a next step, we sought to provide a homogenous synthetic target of uniform size. The lipid phosphatidylserine (PS) is exposed on dying cells and is considered to be an important “eat me” signal (Fadok et al., 1992). PS was previously shown to cause Draper phosphorylation (Tung et al., 2013), but multiple protein ligands for Draper also have been suggested (Kuraishi et al., 2009; Okada et al., 2012). Based upon prior studies, it was unclear whether one or multiple ligands were needed to execute full engulfment. To determine if PS alone might be sufficient to trigger Draper-mediated engulfment, we replaced apoptotic cells with 6.5 μm diameter silica beads coated with a supported lipid bilayer (SLB) containing 10% PS and 0.5% atto390-PE lipid for visualization by microscopy (Fig. 1C). Draper-GFP S2 cells engulfed ∼0.8 PS beads/cell, on average, after a 30 min incubation (Fig. S1A), which was 50-fold higher than the ingestion of beads coated only with phosphatidylcholine (PC) alone. Control GFP transfected S2 cells also engulfed PS-coated beads ∼7 fold less efficiently than the Draper-GFP transfected cells, (Fig. S1A). Draper-GFP S2 cells engulfed beads at 2.5% PS lipid content and showed increasing engulfment up to 12.5% PS (Fig. 1C). The PS concentrations that promote engulfment of beads are within the physiological range present on dead and dying cells (Leventis and Grinstein, 2010). Thus, our S2 cell system is a simplified two-component platform that is dependent on 1) overexpression of the Draper receptor, and 2) PS exposed on a SLB-coated bead.

Next, we examined the activation of Draper in the presence of 10% PS beads. Similar to the apoptotic cell, we found that Shark-mCherry co-localized with Draper-GFP at the interface between PS bead and the Draper-transfected S2 cell (Fig. 1D). A strong pTyr signal was also found at this interface as well (Fig. S1B). Collectively, these results demonstrate that PS ligation is sufficient to trigger Draper locally and transmit the signal from target to cytosol (Fig. 1D), where Draper receptor tails are phosphorylated and recruit the kinase Shark.

We next performed live cell microscopy to visualize the process of Draper-mediated engulfment of PS beads (Fig. S1C, Video 1). We co-transfected Draper-mCherry and LifeAct-GFP in S2 cells and imaged PS bead engulfment by time-lapse imaging. Draper-mCherry expressing cells formed a synapse with PS beads and rapidly began forming a phagocytic cup; F-actin, as assessed by LifeAct-GFP signal, localized at the edges of the cup and the leading edge of the cup extended gradually around the bead. When the phagocytic cup was complete, the PS bead was internalized. Thus, our system is useful for determining the kinetics of PS bead engulfment and following the dynamics of the cytoskeleton and membranes downstream of Draper activation.

### Ligated Draper nucleates mobile microclusters that recruit effector proteins and remodel the actin cytoskeleton

To better visualize the behavior of Draper and the recruitment of proteins to the plasma membrane during the initiation of engulfment signaling, we used Total Internal Reflection Fluorescence-Microscopy (TIRF-M), a technique that confines illumination to within ∼200 nm of the glass coverslip. We coated the glass coverslip with a supported lipid bilayer containing 10% PS (Fig 2A). When Draper-GFP S2 cells settled on this PS supported lipid bilayer, the cells spread rapidly on the surface in a futile effort to engulf the planar surface. Strikingly, Draper-GFP formed microclusters, typically near the leading edge of the cell; these microclusters then underwent retrograde flow from the leading edge towards a central synapse at a rate of ∼1.9 µm/min (n=50) (Fig. 2A; Video 2). The behavior of these Draper clusters is very similar to that reported for adaptive immune receptor microclusters that form in B and T cells and are thought to serve as sites of local signal activation (Kaizuka et al., 2007; Murugesan et al., 2016).

**Figure 2:**
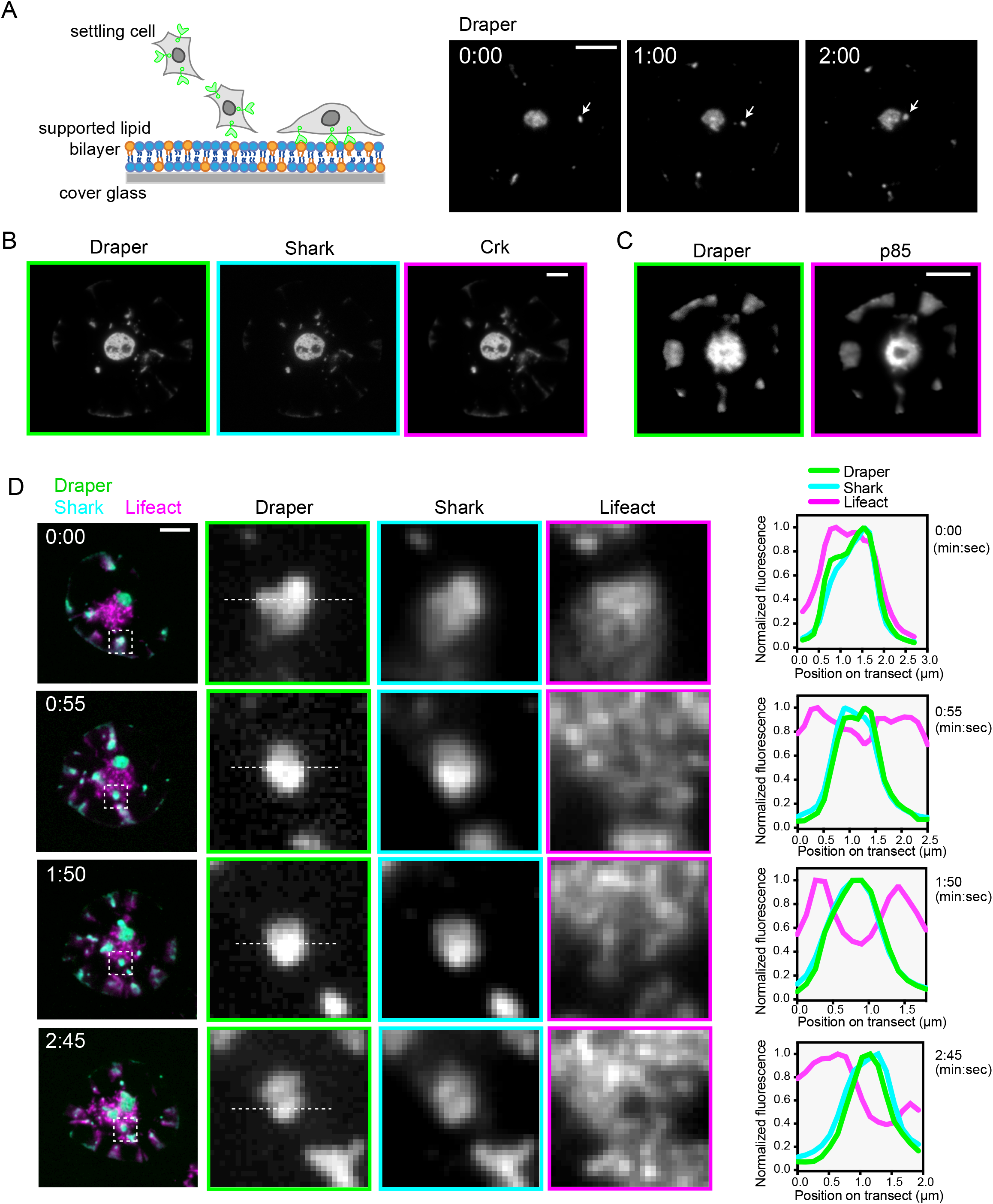
Ligated Draper nucleates mobile microclusters that recruit effector proteins and remodel the actin cytoskeleton. **A.** Schematic of TIRF-M assay setup to visualize ligated Draper-GFP at the plasma membrane. Draper-GFP+ S2 cells were allowed to settle on 10% PS supported lipid bilayer (SLB) prepared on a glass coverslip, and Draper-GFP+ was imaged directly in real time. Draper-GFP forms microclusters on 10% PS SLB. Draper-GFP microclusters visualized by TIRF-M undergo retrograde flow towards a central synapse of Draper-GFP. Microclusters fuse with one-another and are repopulated from the periphery. Time in min:sec. Scale bar, 5 µm. **B.** Draper-GFP microclusters on 10% PS bilayers recruit Shark-TagBFP and Crk-mCherry to the plasma membrane. Cells co-transfected with receptor, kinase and adapter were allowed to settle on 10% PS bilayers and imaged after spreading. Scale bar, 5 µm. **C.** Draper-mCherry microclusters on 10% PS bilayers recruit the activating subunit of PI3K (p85-GFP) to the plasma membrane. Cells co-transfected with Draper-mCherry and p85-GFP were allowed to settle on 10% PS bilayers as above and imaged after spreading. Scale bar, 5 µm. **D.** Draper-GFP microclusters on 10% PS bilayers recruit Shark-mCherry but exclude the F-actin reporter Lifeact-TagBFP in a time-dependent manner. Cells co-transfected with Draper-GFP and Lifeact-TagBFP were allowed to settle on 10% PS bilayers as above and imaged at 55 sec intervals after spreading. The white box highlights the expanded region at right to demonstrate that peripheral Draper-GFP microclusters co-localize with Shark-mCherry and Lifeact-TagBFP initially and appear to locally deplete F-actin over time, as indicated by reduced Lifeact-TagBFP signal as the cluster matures. Transect showing position of plotted normalized fluorescence is indicated as a dashed line. Time in min:sec. Scale bar, 5 µm.

Next we asked if Draper-GFP clusters are sites of recruitment for downstream signaling molecules. Draper is proposed to interact with Shark, a dual SH2 protein, and work with the adapter protein Crk, which harbors one SH2 and two SH3 domains. To test if these proteins are recruited to Draper microclusters, we allowed Draper-GFP, Shark-TagBFP, and Crk-mCherry transfected S2 cells to settle on 10% PS bilayers and imaged them by TIRF microscopy. Activated Draper co-localized with Shark kinase, the adapter Crk, and the activating subunit of PI3K, p85 (Fig. 2B, Fig. 2C, Fig. S2, Video 3). Cytosolic phosphotyrosine binding proteins co-localized with Draper at peripheral microclusters immediately upon formation and persisted during migration toward the large central synapse (Fig. 2B, Fig. 2C). Collectively, these results demonstrate that upon ligation to PS, Draper is phosphorylated and nucleates formation of signaling microclusters, mobile assemblies that recruit multiple SH2-domain containing proteins.

To determine if Draper clusters have local effects on the actin cytoskeleton, we co-expressed Draper-mCherry, Shark-GFP, and LifeAct-BFP and allowed these cells to settle on PS bilayers. The appearance of Draper-mCherry microclusters was accompanied by an almost simultaneous colocalization signal with SharkGFP and LifeAct-BFP (Figure 2D; first time point, 0:00). Surprisingly, over time and while undergoing retrograde transport, the Shark-GFP signal remained consistently strong, while the LifeAct-BFP signal became depleted under the microcluster (Figure 2D). These results suggest that Draper signaling temporally downstream of Shark recruitment leads to highly localized depolymerization of the actin cytoskeleton. Remarkably, even though locally depleted of filamentous actin, the Draper signaling clusters remain well coupled to the actin retrograde flow overall and were transported to the cell center.

### Exclusion of a transmembrane phosphatase at the ligated membrane-membrane interface

We next used our reconstitution system to define how PS triggers Draper phosphorylation. Both the T cell receptor (TCR) and Fc receptors are believed to be triggered by the exclusion of the transmembrane phosphatase CD45, which has a bulky extracellular domain, from zones of close membrane-membrane contact created by the pMHC-TCR or IgG-FcR binding (Choudhuri et al., 2005; Burroughs et al., 2006; James and Vale, 2012; Freeman et al., 2016). For TCR triggering, the Src family kinase Lck is attached to the membrane inner leaflet and remains uniformly distributed after pMHC-TCR engagement, so phosphatase exclusion shifts the kinase-phosphatase balance towards net receptor phosphorylation. In contrast, EGFR and GPCRs are triggered via receptor conformational change (Dawson et al., 2005; Erlandson et al., 2018). We sought to determine which mechanism governs Draper triggering. The segregation model makes two key predictions: 1) a phosphatase with a bulky extracellular domain should be excluded from phosphorylated receptor microclusters and 2) bringing the target membrane and phagocyte membrane into close proximity via a heterologous interaction module should trigger receptor phosphorylation and signaling. We sought to test whether a segregation mechanism might function to trigger Draper phosphorylation. Drosophila S2 cells express several transmembrane phosphatases as measured by RNA-Seq (Zhang et al., 2010) (Fig. 3A). We tested the transmembrane phosphatase Ptp69D, because of its high expression level and similar domain structure to the segregated mammalian phosphatase CD45 (extracellular fibronectin repeats and cytosolic tyrosine phosphatase activity (Fig. 3A).

**Figure 3:**
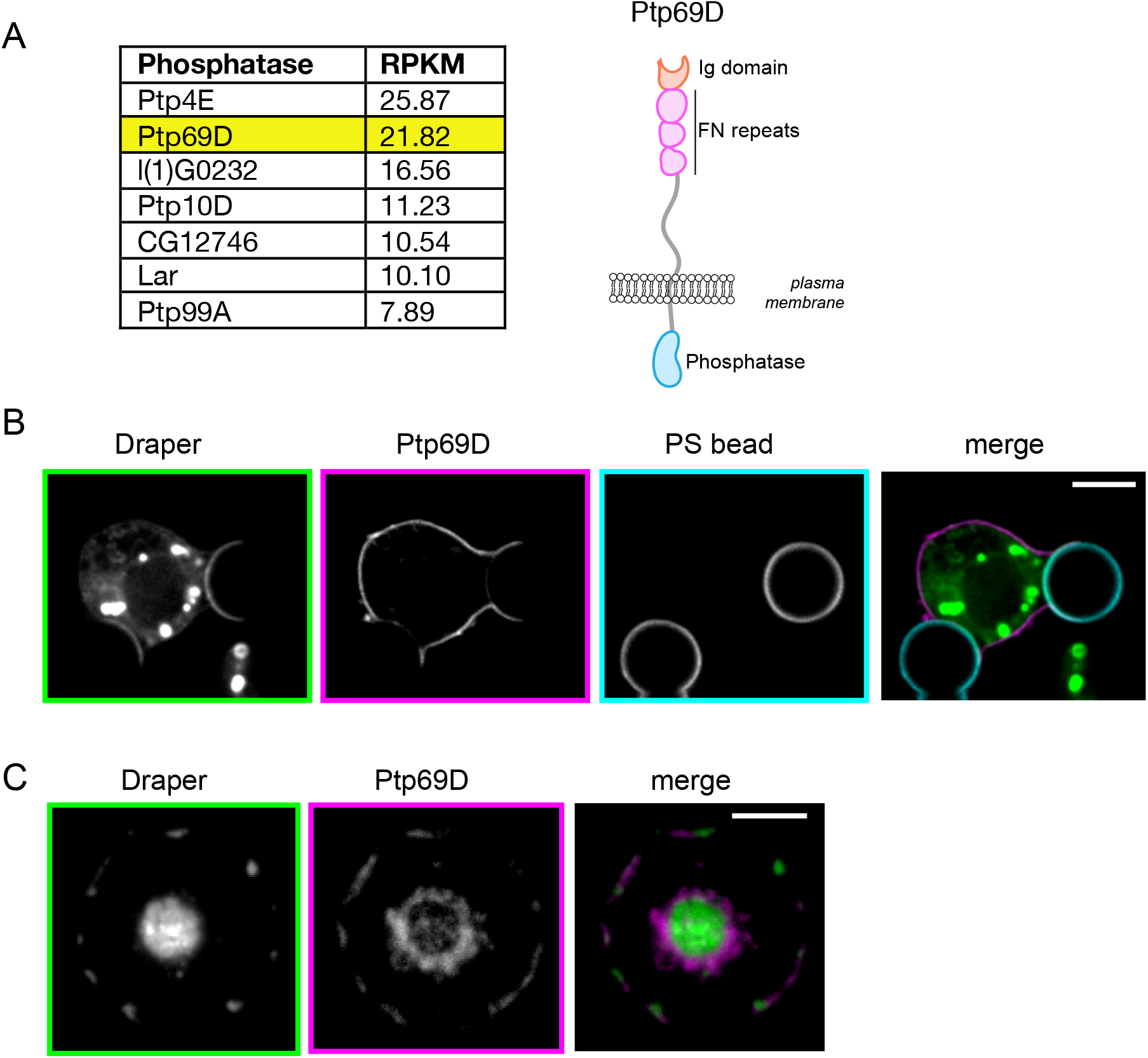
Exclusion of a transmembrane phosphatase at the ligated membrane-membrane interface. **A.** S2 transmembrane phosphatase expression data and schematic of Ptp69D-GFP domain structure, highlighting features of its bulky extracellular domain. The transmembrane phosphatase Ptp69D segregates away from Draper at synapses between 10% PS beads and the S2 cell. Draper-mCherry and Ptp69D-GFP were co-transfected into S2 cells and localization at the synapse assessed after a 15 min incubation. **B.** Draper-mCherry clusters exclude Ptp69D-GFP on 10% PS supported lipid bilayers (SLBs). Draper-mCherry/Ptp69D-GFP expressing S2 cells were allowed to settle and spread on bilayers and localization of receptor and phosphatase assessed after 15 minutes. Phosphatase exclusion was observed both at peripheral clusters and the central synapse. Scale bar indicates 5 µm.

To determine if Ptp69D segregates away from Draper at the ligated interface, we co-transfected the two proteins with different fluorescent protein tags and assessed localization in living cells. Microscopy revealed that ligated Draper-mCherry concentrated at the synapse between the S2 cell and PS bead, while Ptp69D-GFP was partially excluded from this zone (Fig. 3B). Draper microclusters that formed at the interface between an S2 cell and a planar PS-containing supported lipid bilayer also excluded Ptp69D in TIRF (Fig. 3C). Thus, Ptp69D segregates away from ligated Draper, a result that is consistent with a kinetic segregation triggering mechanism.

### A synthetic receptor promotes Draper-dependent engulfment and actin remodeling

In the kinetic segregation model, two opposing membranes are brought into close apposition by the binding energy between receptor and its ligand, which results in the exclusion of larger transmembrane phosphatases (Cordoba et al., 2013; James and Vale, 2012; Carbone et al., 2017). To further examine whether Draper may trigger phosphorylation through a kinetic segregation mechanism rather than a receptor conformational change, we tested whether the extracellular domain of Draper and its ligand PS could be replaced with an artificial receptor-ligand pair. Previous studies on T cell receptor triggering showed that the extracellular domain of the TCR and pMHC could be substituted with FKBP and FRB, which upon addition of rapamycin to bridge FKBP and FRB, trigger TCR phosphorylation (James and Vale, 2012). We created a similar inducible system for Draper by replacing its endogenous extracellular domain with FRB (FRB-EXT-Draper-INT) and replacing PS with His_10_-FKBP, which was bound to the SLB on the bead via a NiNTA-lipid (Fig 4A). In the absence of rapamycin, beads were engulfed at very low levels (0.003 beads/cell). However, in the presence of rapamycin, the engineered receptor promoted robust engulfment of FKBP-bearing beads (Fig. 4A, Video 5). Live cell microscopy revealed that the FRB-EXT-Draper-INT concentrated at the cell-bead interface and recruited Shark kinase, suggesting that the chimeric receptor, like Draper, is activated by tyrosine phosphorylation (Fig 4B). Like Draper-GFP, FRB-EXT-Draper-INT excluded the phosphatase Ptp69D-GFP (Fig. 4C), consistent with a kinetic segregation-based triggering mechanism. Thus a chimeric receptor that is designed to interact with a non-physiological ligand is functional for local receptor triggering, phosphatase exclusion, and engulfment.

**Figure 4:**
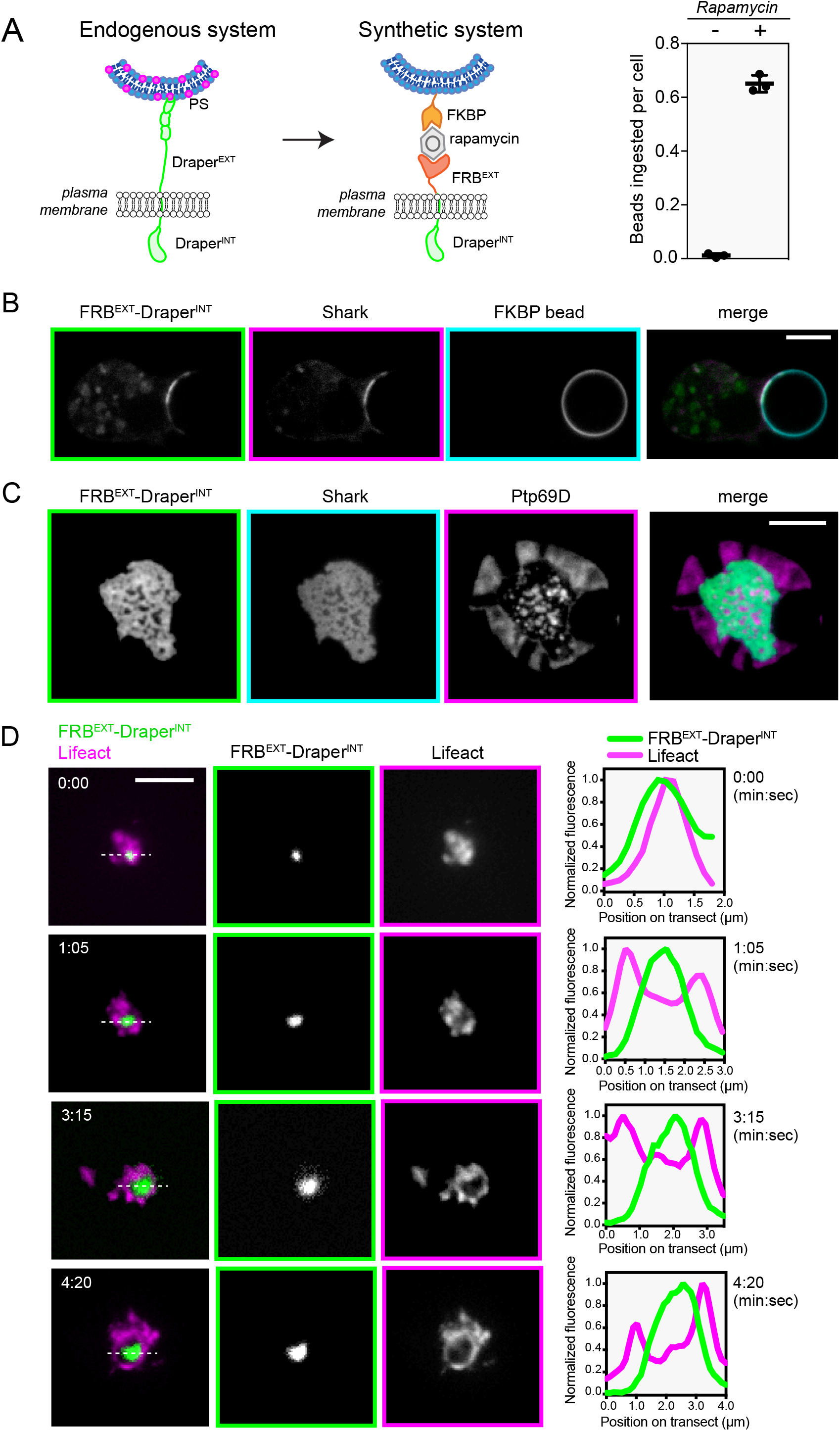
A synthetic receptor promotes Draper-dependent engulfment and actin remodeling. **A.** Schematic showing extracellular FRB fused to the Draper transmembrane and cytosolic domains (FRB-EXT-Draper-INT) to program clearance signaling towards FKBP-bearing beads (see Methods). FRB and FKBP are brought together into a 100 fM ternary complex in the presence of 1 µM rapamycin. FRB-EXT-Draper-INT promotes specific phagocytosis of FKBP beads. S2 cells transfected with FRB-EXT-Draper-INT were incubated with 10 nM FKBP beads for 30 min, after which bead ingestion was quantified after imaging using spinning disk confocal microscopy (see Methods). The mean number of beads fully ingested per cell for three independent biological replicates are shown (mean ± SD). Cells were incubated either with 1 µM rapamycin (right, +) or an equivalent volume of DMSO vehicle (left, -). **B.** The kinase Shark is recruited to FRB-EXT-Draper-INT, indicating local receptor activation by tyrosine phosphorylation. S2 cells expressing FRB-EXT-Draper-INT-GFP and Shark-Cherry were incubated with FKBP beads coupled to 10 nM his10 protein (see Methods) in the presence of 1 µM rapamycin for 15 min and localization was assessed. A middle section from a confocal z-stack is shown. **C.** S2 cells were transfected with FRB-EXT-Draper-INT-mCherry, Shark-TagBFP, and Ptp69D-GFP and allowed to settle on SLB with bound His_10_-FKBP bilayers in the presence of 1 µM rapamycin for 15 min. Ptp69D segregates away from activated Draper at the interface between FKBP bilayer and S2 cell. In contrast, Draper and Shark overlapped, indicating a zone of local receptor activation and phosphorylation. Scale bars, 5 µm. **D.** FRB-EXT-Draper-INT microclusters on FKBP bilayers exclude the F-actin reporter Lifeact. Cells co-transfected with FRB-EXT-Draper-INT-mCherry and Lifeact-GFP were allowed to settle on His_10_-FKBP bilayers as above and imaged at 65 sec intervals. FRB-EXT-Draper-INT-mCherry microclusters co-localize with Lifeact-GFP initially and appear to locally deplete F-actin, as indicated by reduced Lifeact-GFP signal as the cluster matures. Transect showing position of plotted normalized fluorescence is indicated as a dashed line. Time in min:sec. Scale bar indicates 5 µm.

Using Lifeact-GFP-expressing cells, we also examined whether microclusters nucleated by the FRB-EXT-Draper-INT receptor locally deplete F-actin. Like Draper, the FRB-EXT-Draper-INT receptor and Lifeact colocalized in the TIRF field, but over 1 min, the F-actin signal became depleted under the microcluster, creating the appearance of a small actin hole (Figure 4D, Video 6). Collectively, our results using the engineered FRB-EXT-Draper-INT receptor indicate that formation of a ligated zone that excludes a transmembrane phosphatase is sufficient to trigger receptor phosphorylation, effector recruitment, and local F-actin depletion observed for Draper.

### Ordered multisite Tyr phosphorylation is required for full Draper activation

We first sought to determine which Tyr residues are essential for Draper-dependent engulfment in our cellular reconstitution assay. The Draper tail domain contains an ITAM motif (YxxI/Lx_6-12_YxxI/L), an NPxY motif thought to interact with CED-6 (Fujita et al., 2012), and 10 other tyrosine residues, 2 of which are in close proximity to the transmembrane domain. First, we created a construct in which 11 of the 13 tyrosines were mutated to alanine (Draper-Ala11-GFP), leaving the two tyrosines that are immediately adjacent to the transmembrane segment intact since they potentially could have a structural role. We then compared phagocytic proficiency of cells transfected with Draper-Ala11-GFP mutant to full-length Draper-GFP and GFP alone (endogenous background phagocytosis). Consistent with a role for Tyr phosphorylation on Draper to promote phagocytosis, Draper-Ala11-GFP ingested PS-beads at levels that were comparable to the GFP control and ∼15-fold below Draper-GFP cells after a 45 min incubation (Fig 5A).

**Figure 5:**
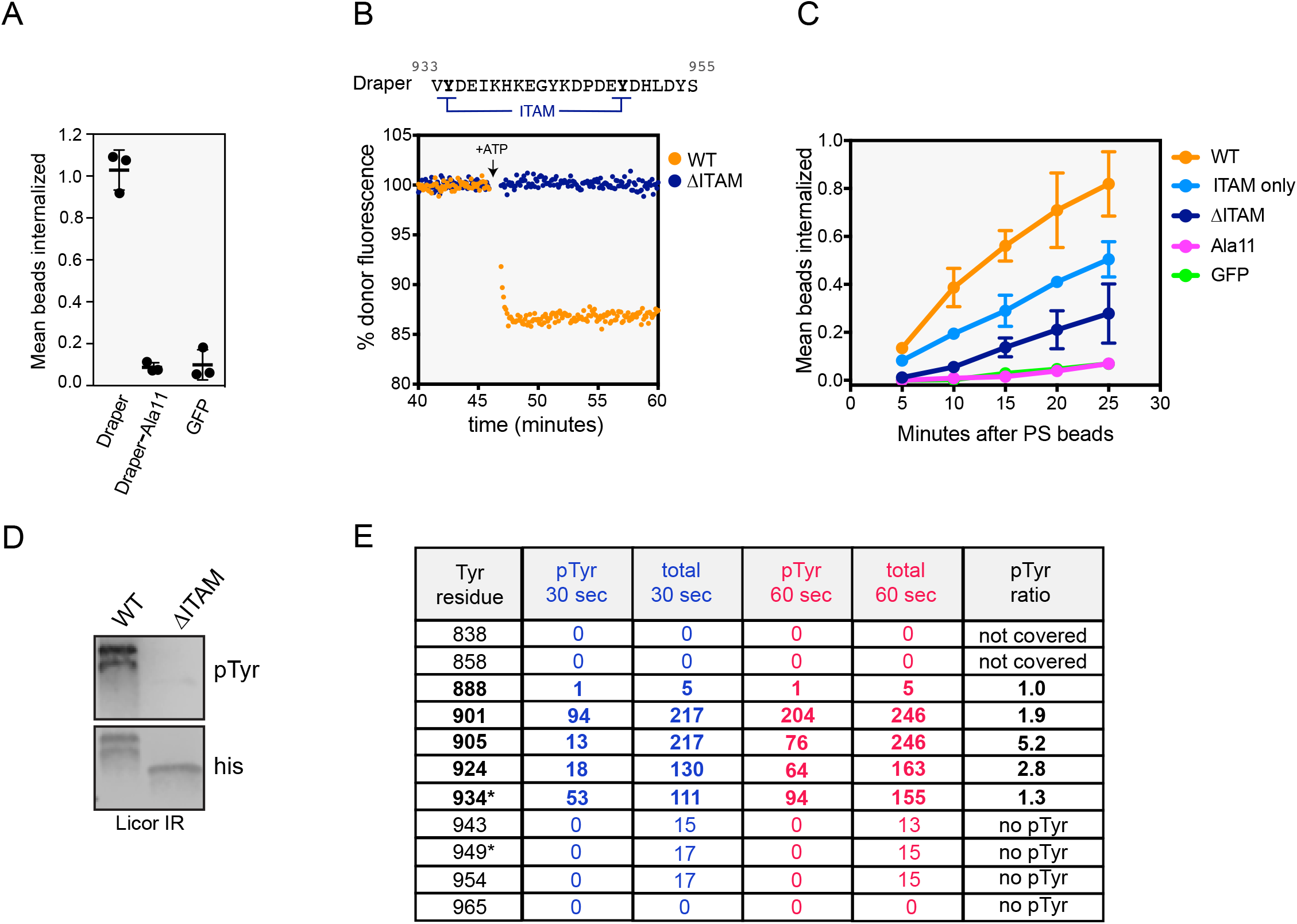
Ordered multisite Tyr phosphorylation is required for full Draper activation. **A.** Engulfment of 10% PS beads by S2 cells expressing Draper-GFP, Draper-Ala11-GFP, or a GFP control was assessed after 45 min incubation. Mean number of beads ingested per cell for three biological replicates are shown (mean ± SD). **B.** Alignment of residues mutated to Phe in Draper-ΔITAM. When a BG505-labeled reporter protein comes into close proximity to a rhodamine labeled liposome, the 505 nm fluorescence signal is quenched. If quenching occurs after addition of ATP it is most likely due to the Src42a-dependent phosphorylation of Draper’s cytoplasmic tyrosine residues and effector recruitment. The results shown here reveal that direct binding of BG505-Shark-tSH2 to Draper’s tail is dependent on an intact ITAM on Draper. His_10_-Draper cytosolic domain or His_10_-DraperY934F/Y949F (ΔITAM) receptor tail were ligated to 0.5% rhodamine-PE liposomes and ATP dependent quenching was assessed. **C.** Draper-ΔITAM displays a kinetic delay and reduced phagocytic proficiency relative to Draper-GFP, while Draper’s ITAM (ITAM only) is sufficient for partial engulfment activity. S2 cells transfected with Draper-GFP or indicated mutants were incubated with 10% PS beads, allowed to settle, and imaged at the indicated time points. The mean number of beads fully ingested per cell for three independent biological replicates are shown (mean ± SD). **D.** On-liposome phosphorylation reactions to assess Draper activation by its cognate Src family kinase Src42a. His_10_-Draper cytoplasmic domain or 10his-DraperY934F/Y949F (ΔITAM) receptor tail (both at 1 µM) were ligated to liposomes doped with DGS-Ni-NTA and incubated with 86 nM Src42a and 1 mM ATP for 30 min. Reactions were quenched with SDS-PAGE buffer containing DTT and 2-Me and boiled for 10 min at 95°C. Samples were immunoblotted with phosphotyrosine antibody or His_10_ antibody. Quantification of phosphorylation of WT or ΔITAM by Src42a was determined as the ratio of total pTyr signal over total His_10_ signal internally for each lane. **E.** 1 µM His_10_-Draper cytoplasmic domain was phosphorylated using 1 nM on-membrane Src42a and samples quenched using 10 mM EDTA and 8 M Urea. Reactions were digested for 2D mass spectrometry at 30 second and 60 second timepoints (see Methods)**. E.** 2D mass spectrometry peptide counts for each Tyr residue on Draper’s cytoplasmic tail. pTyr and total peptide counts for each timepoint are shown, 30 sec in blue, 60 sec in red. pTyr in the right column indicates the ratio of (pTyr peptides at 60 sec/all peptides) relative to the (pTyr peptides at 30 sec/all peptides). A ratio above 1.0 indicates preferential phosphorylation at the later 60 sec timepoint. “Not covered” indicates no peptides were detected for a residue in the 2D mass spec. “No pTyr” indicates that no peptides with the +79.9663 Da shift characteristic of phosphorylation were detected.

We next sought to dissect the roles of Draper’s cytoplasmic Tyr residues directly using a purified biochemical system on liposomes. The cytoplasmic tail of Draper and its cognate Src family Drosophila kinase Src42a (Ziegenfuss et al., 2008) were purified and bound via a His10 tag to NiNTA-modified lipids in the liposome. To detect binding between Shark and Draper, we used a fluorescently-labeled tandem SH2 Shark construct (BG505-tSH2-Shark) whose signal quenches upon binding to Draper as it comes in proximity to a lipid-bound acceptor dye (Hui and Vale, 2014). After ATP was added as substrate for Src42a-mediated phosphorylation, we observed a time-dependent quenching of Shark fluorescence as it bound to the phosphorylated receptor (Fig. 5B, orange trace). Next, we tested the Draper receptor with double tyrosine to phenylalanine mutation in its ITAM motif (Fig. 5B, Draper-ΔITAM). Draper-ΔITAM showed little or no recruitment of Shark (Fig. 5B, blue trace). This biochemical result confirms that Shark binds Draper directly via Draper’s ITAM (Ziegenfuss et al., 2008).

We next tested the role of the Draper ITAM in our cell-based system by mutating its two Tyr residues to Phe (Draper-ΔITAM; residues in 5B) and assessed the ability of this mutant to promote phagocytosis. Draper-ΔITAM showed a kinetic delay and a lower number of beads ingested (Fig. 5C) compared to wildtype Draper, revealing its critical role in signaling and Shark recruitment (Fig. S4). However, Draper-ΔITAM still promoted phagocytosis to a greater extent than the effectively null Draper-Ala11 construct (Fig. 5C).

We also tested if Draper’s ITAM is sufficient to signal by adding back the ITAM tyrosines (Y934 and Y949) to the null Ala-11 mutant (Draper ITAM-Only). The Draper ITAM-Only receptor promoted phagocytosis of PS beads, but did so more slowly than wildtype and to a lesser extent than wildtype Draper (Fig. 5C). As expected, the Draper ITAM was necessary and sufficient to recruit Shark kinase to the synapse between S2 cell and PS bead (Fig. S4). Collectively, our mutational analyses suggest that Draper’s ITAM plays a critical role in signaling but that non-ITAM residues also contribute to full receptor activity during engulfment.

To further understand the mechanism of Src42a-dependent activation of Draper, we tested the extent of Tyr phosphorylation on Draper’s cytoplasmic tail in the on-membrane biochemical assay. After a 30 min kinase reaction, immunoblotting with anti-phosphotyrosine antibodies revealed a noticeable shift in the electrophoretic mobility of Draper, which is suggestive of phosphorylation on multiple Tyr residues (Fig. 5D). Remarkably, even though the Draper-ΔITAM contains 9 additional tyrosine residues, its phosphorylation as detected by phosphotyrosine immunoblotting was minimal (5% of wildtype Draper) in the same kinase reaction (Fig. 5D). We observed a similar result using the Src family kinase Lck (Fig. S5), suggesting that preferential modification through ITAM residues is a general mechanism of Draper receptor activation.

Greatly diminished phosphorylation of the Draper-ΔITAM mutant could arise because 1) only the ITAM residues are phosphorylated by Src family kinases, or 2) phosphorylation of the ITAM motif is needed for the efficient phosphorylation of non-ITAM tyrosines in Draper’s tail. To distinguish between these possibilities, we repeated the Draper phosphorylation assay and quenched reactions at 30 and 60 sec after addition of ATP. To determine if any residues on Draper are phosphorylated in ordered fashion, we performed 2D mass spectrometry on the 30 and 60 sec samples. This method yielded peptides covering Tyr residues on Draper and allowed us to differentiate between phosphorylated and non-phosphorylated residues (Fig. 5E). By comparing the proportion of phosphorylated peptides for each site at both 30 and 60 sec, we could determine whether certain Tyr residues on Draper are preferentially modified over time. The ITAM Y934 residue was modified to a similar extent at both time points (pTyr ratio of 1.3, Fig. 5E). However, the non-ITAM residues (except for Y888 which also had few phosphopeptides detected) were preferentially phosphorylated later, as indicated by the 60 sec/30 sec ratios ranging from 1.9 to 5.2 (Fig. 5E). Collectively these results indicate that multiple Draper tyrosines are phosphorylated by Src42a, and that phosphorylation of non-ITAM residues strongly depends upon prior phosphorylation of the ITAM residues.

## Discussion

In this study, we converted poor phagocytes (Drosophila S2 cells) into proficient eaters by expressing a single receptor, Draper. We also show that PS is sufficient to locally trigger receptor phosphorylation and engulfment signaling, building on previous work that demonstrated PS-liposome dependent Draper phosphorylation (Tung et al., 2013). Thus, we developed a simplified assay platform to dissect a complex cell signaling process that requires multiple receptors and ligands in vivo (Freeman and Grinstein, 2014). Draper is also reported to recognize the protein ligands Pretaporter and DmCaBP1 (Kuraishi et al., 2009; Okada et al., 2012) and has been reported to work in collaboration with co-receptors in flies such as Six Microns Under (SIMU) (Kurant et al., 2008) and integrins (Nagaosa et al., 2011). Our results do not rule out a role for these proteins in vivo to tune sensitivity for triggering and/or allow engulfment under low concentrations of PS (Manaka et al., 2004). However, our study shows that Draper interaction with PS provides a minimal system for studying apoptotic cell engulfment.

Using this cellular platform, in conjunction with a liposome-based biochemical assay, we have been able to gain insight into the initial steps of the corpse clearance signaling pathway (see model Fig. 6). Our results show that ligation to the lipid PS is sufficient to promote Draper clustering and segregation of a transmembrane phosphatase at the ligated interface. Spatial separation between ligated Draper clusters and transmembrane phosphatases shifts a kinase-phosphatase balance in favor of receptor phosphorylation by Src42a. Phosphorylation of Draper’s tail activates a downstream cytosolic signaling cascade, beginning with the recruitment of Shark and other phosphotyrosine binding effector proteins, to Draper microclusters. After effector recruitment, early steps of Draper signaling include localized depolymerization of the actin cytoskeleton. Further downstream signaling likely leads to integrin activation and other events that lead to the formation of the phagocytic cup, although these more distal events in the pathway were not the focus of this study.

**Figure 6:**
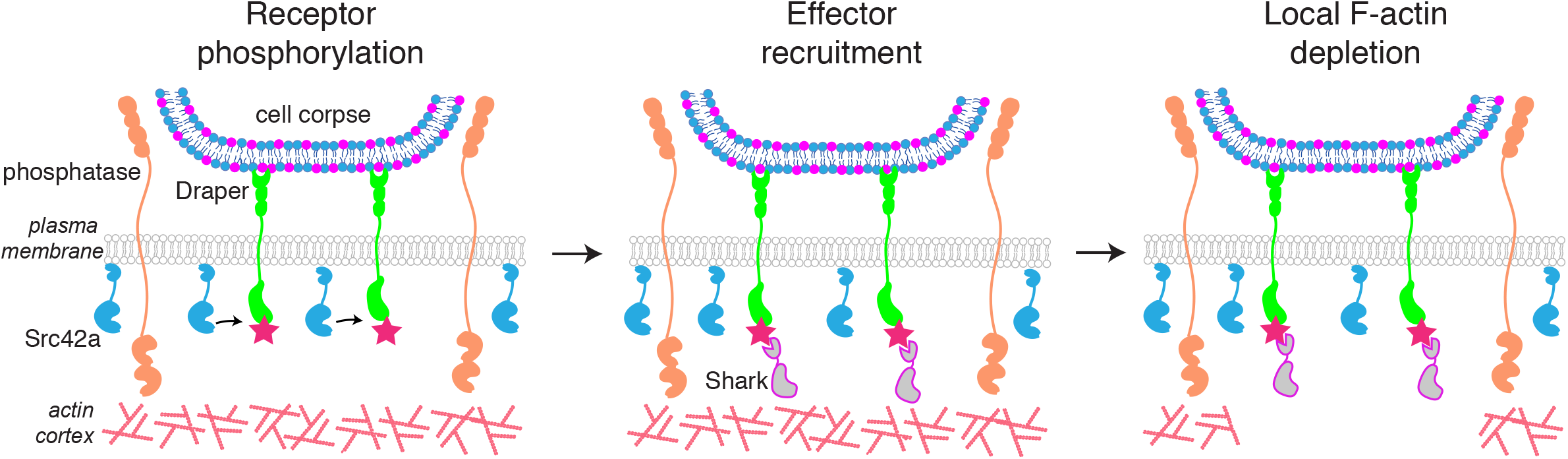
A mechanism for engulfment signaling initiation. Schematic representation of the early molecular steps of corpse clearance through the Draper receptor. *Left:* Draper ligation to PS on a cell corpse creates an exclusion zone where ligated receptors cluster at the synapse and unligated molecules such as bulky transmembrane phosphatases segregate away from receptors. Spatial segregation between receptors and inhibitory phosphatases shifts the balance towards receptor phosphorylation by Src42a. *Middle:* Tyr phosphorylation on Draper’s tail recruits cytosolic effector proteins including the kinase Shark. *Right:* Effector recruitment promotes ordered signaling outputs including local depletion of F-actin at ligated microclusters.

### Comparison between Draper signaling and mammalian immune signaling

A potential similarity between Draper signaling and mammalian immune receptor activation was first suggested by the finding of an ITAM in the Draper cytoplasmic tail and the ability of this ITAM to bind Shark kinase in yeast two-hybrid and pull-down assays (Ziegenfuss et al., 2008). Our findings extend this work by providing evidence that Draper, like the T cell and Fc immune receptors, is triggered by a kinetic segregation mechanism. The kinetic segregation model, initially proposed as a mechanism for T cell activation, postulates that apposition of a membrane-membrane interface at the synapse between the T cell and antigen presenting cell (APC) results in the exclusion of a bulky transmembrane phosphatase CD45, which shifts a kinase-phosphatase equilibrium in favor of receptor phosphorylation (Davis and van der Merwe, 2006). A segregation mechanism was recently proposed to regulate FcR activation, another mammalian immune signaling pathway, by creating an integrin-based diffusion barrier that excludes CD45 (Freeman et al., 2016).

Prior to our work, it remained unclear if ligated corpse clearance receptors also use phosphatase exclusion to create zones of receptor triggering. Here, we show that Draper ligation to PS on a second membrane excludes a transmembrane phosphatase (Ptp69D) from the synapse at the region of receptor phosphorylation and effector recruitment. Similar results showing exclusion were obtained with our chimeric receptor based upon FKBP-rapamycin-FRB interaction. In the case of T cells, a single transmembrane phosphatase (CD45) is highly expressed on the surface and appears to be the dominant phosphatase in regulating signaling. However, phagocytic cells express several transmembrane phosphatases, including Ptp69D, which together may contribute to regulating Draper and other phagocytic receptors. In the future, it will be illuminating to determine how the specific composition of phosphatases on the membrane (different sizes, localization patterns, and expression levels) are used to tune the sensitivity of engulfment. It also will be interesting to explore whether Draper, like FcR (Freeman et al., 2016), activates integrin signaling to create a diffusion barrier for CD45.

Draper ligation to PS results in the formation of receptor microclusters, physical assemblies that organize engulfment effectors. Draper microclusters are transported by the actin cytoskeleton from the periphery to the cell center, a behavior very similar to that described for the TCR (Bunnell et al., 2002; Douglass and Vale, 2005; Varma et al., 2006). The striking similarity extends to the recruitment of key signaling molecules to the microclusters. We show that, similar to the TCR, Draper recruits several SH2-domain containing effector proteins, including Shark kinase, the adapter Crk, and the PI3K subunit p85.

While the formation and retrograde transport of microclusters appears to be similar for the TCR and Draper, we observed a striking difference with regard to the effects of these clusters on filamentous actin (F-actin). The TCR microclusters reside above actin-rich zones as they are transported from the periphery to the central synapse (cSMAC) (DeMond et al., 2008; Kaizuka et al., 2007; Murugesan et al., 2016; Yi et al., 2012). We also observed that Draper microclusters initially co-localize with the F-actin reporter Lifeact. However, as the clusters mature, but well before they reach the central synapse, the microclusters deplete F-actin to create “holes” in the actin cytoskeleton. We observed a similar anti-correlation between Draper and F-actin at microclusters of the synthetic FRB-EXT-Draper-INT receptor (Fig. 4D) and at the large synapse between Draper-expressing S2 cell and PS bead (Fig. S3). Our findings indicate that F-actin depletion over time both at peripheral microclusters and the central synapse is a distinct output of Draper receptor activation to actin regulators compared to the TCR signaling pathway.

Actin-depleted zones have been observed to form at the cSMAC at the mature synapse between a T cell and its target (Dustin, 2014; Ritter et al., 2015). Ritter et al. proposed that signals from the microtubule cytoskeleton might clear the actin cytoskeleton in preparation for cytolytic granule secretion. For Fc receptor-mediated engulfment, actin-depleted zones have been reported at the interface between the phagocyte and a bead (Schlam et al., 2015; Yamauchi et al., 2012); actin depletion was suggested to be important for the phagocytosis of large but not small targets (Schlam et al., 2015). In addition, elegant recent work by Freeman et al. (2016) reported that macrophages interacting with micropatterned, immobilized IgG will polymerize F-actin surrounding the ligated Fc receptors, thereby forming “actin rings”. This study focused on integrin-mediated actin assembly at the periphery outside of the zone of receptor-ligand engagement, but images in Freeman et al. (2016) also reveal actin-depletion at the center of the receptor zones in the micropatterned array.

The kinetics of actin depletion observed in this study suggests that actin depletion is an active part of the Draper signaling pathway that follows receptor phosphorylation by ∼1 min and is restricted to the zone of ligated receptors. The molecular mechanism responsible for actin disassembly could involve: 1) a repression of Arp2/3 actin nucleation near the phosphorylated Draper receptor, potentially in combination with a barrier that presents nearby actin filaments from polymerizing or moving into the receptor zone, 2) RhoGAP repression of Rac and Cdc42, as suggested by Schlam et al. for large phagocytic cups (2015), or 3) the recruitment and/or activation of actin depolymerizing factors, such as cofilin, at the Draper signaling zone. Additional work will be needed to distinguish between these models.

### Mechanism of Draper triggering upon ligation

The current model for T cell activation and FcR-mediated phagocytosis suggests that triggering is driven entirely by phosphorylation of ITAM tyrosines. In those cases, there are not additional tyrosine residues outside of the ITAM motifs. Shark kinase, like ZAP70 and Syk, binds specifically to the ITAM motif on Draper and we show that ITAM phosphorylation plays an important role in engulfment signaling. However, unlike the TCR and FcR, we show that Draper contains non-ITAM tyrosines that are also phosphorylated during activation and are functionally important for engulfment. We find that non-ITAM tyrosines play a role in Draper activation, thus supporting a signaling function both for Draper’s ITAM and non-ITAM residues as was suggested earlier (Fujita et al., 2012; Ziegenfuss et al., 2008). By measuring Draper tyrosine phosphorylation directly via mass spectrometry we report modification of residues outside of Draper’s ITAM. Importantly, Draper’s non-ITAM tyrosine residues contribute to engulfment efficiency in cell-based assays. The phosphorylated non-ITAM tyrosines in Draper may bind additional SH2-containing adapter proteins or other effectors. Further studies will be needed to identify additional direct binding partners of the phosphorylated cytoplasmic tail domain of Draper and determine the higher order structure of signaling microclusters that promote engulfment signaling.

Our biochemical reconstitution also reveals that Draper’s ITAM domain is required for efficient phosphorylation of the non-ITAM tyrosines. This result indicates a sequential mechanism for multisite phosphorylation on Draper. One model for how this might occur is through the direct recruitment of Src family kinases to the phosphorylated ITAM domain of Draper. We show that the Drosophila Src42a kinase has a higher specificity for the tyrosines in Draper’s ITAM domain and phosphorylates these residues first. We also confirmed ITAM-specificity for Draper using Lck, another Src family kinase that phosphorylates ITAM domains on TCR cytosolic tails (Fig. S5). One possible mechanism for the sequential phosphorylation involves a mechanism in which a kinase creates its own binding site before modifying other residues. This idea has been suggested for Src-dependent phosphorylation of synthetic ITAM-containing substrate peptides (Pellicena et al., 1998) and phosphorylation of p130^CAS^ by Abl kinase (Mayer et al., 1995). A similar phenomenon was observed for Lck-dependent modification of sequential ITAM residues on the T cell receptor, which is required for full activation of this receptor (Lewis et al., 1997). Thus, our data could be explained by Src42a binding to the phosphorylated ITAM through its own SH2 domain and then phosphorylating non-ITAM residues on Draper (Fig. 5). Thus, in common with other signaling systems, Draper phosphorylation involves a hierarchal phosphorylation of tyrosines in its tail domain and this ordered sequence of phosphorylation events is required for full receptor activation.

### Synthetic receptors to redirect engulfment signaling

We show here that the extracellular Draper-PS receptor-ligand pair can be functionally replaced by a synthetic module, FKBP-rapamycin-FRB. The ability of this synthetic receptor to promote engulfment is analogous to findings for the T cell receptor (TCR), which can be triggered using the same extracellular FKBP-rapamycin-FRB module (James and Vale, 2012). These synthetic receptors create signaling zones for both Draper (Fig. 3E) and TCR (James and Vale, 2012) that exclude a transmembrane phosphatase to favor phosphorylation of intracellular receptor signaling domains. These programmed receptors enable activation of intracellular signaling pathways independently of their native extracellular domains. Therefore, replacing Draper and PS with a synthetic FKBP-rapamycin-FRB module provides a mechanistic basis to reprogram engulfment signaling toward new targets. Synthetic engulfment receptors consisting of intracellular receptor signaling tails fused to non-native extracellular domains is conceptually similar to chimeric T cell receptors (CAR-Ts) that recognize cancer antigen to promote specific cell killing (Lim and June, 2017). A mechanistic basis to program engulfment signaling indicates that in the future engulfment signaling could be directed towards functionally relevant targets such as cancer cells or pathogens.

## Acknowledgments

We thank N. Stuurman for help with microscopy, C. Carbone, E. Hui, N. Kern, M. Morrissey, X. Su, and M. Taylor for reagents and discussions, and L. Kohlstaedt and Y-S Kim (Vincent J. Coates Proteomics/Mass Spectrometry Laboratory at UC Berkeley, supported in part by NIH S10 Instrumentation Grant S10RR025622) for performing the mass spectrometry. We also thank K. McKinley, M. Morrissey, E. Hui, A. Jain, X. Su, T. Skokan, and P. Williamson for helpful critiques of the manuscript. A.P.W. was supported by a CRI-Irvington Postdoctoral Fellowship. This work was funded by the Howard Hughes Medical Institute and the National Institutes of Health (5R35GM118106-02, R.D.V.).

## Author Contributions

A.P.W. and R.D.V. conceived of the study and designed the experiments. A.P.W. performed the experiments, analyzed the data, and prepared the figures. A.P.W. and R.D.V. wrote the manuscript.

## Video Legends

**Video 1: Cellular reconstitution of Draper signaling**

Video of stills shown in Fig. S1C. An S2 cell co-transfected with Draper-GFP and Lifeact-BFP was incubated with 10% PS/0.5% atto647-PE 6.4 µm silica beads. Video is a maximum projection of a z-stack (1 µm between z-positions). Z-stacks captured by spinning disk confocal microscopy were recorded every 3 min. Time in minutes. Scale bar, 5 µm.

**Video 2: Formation of mobile Draper-GFP microclusters on 10% PS bilayers**

Video of stills shown in Fig. 2A. An S2 cell transfected with Draper-GFP was allowed to settle on 10% PS supported lipid bilayers and imaged by TIRF microscopy. Cell was imaged every 15 sec. Draper-GFP forms mobile microclusters that appear at the edge of the spread cell and migrate toward the prominent central synapse of Draper-GFP signal. Time in min:sec. Scale bar, 5 µm.

**Video 3: Visualization of signaling microclusters on 10% PS bilayers**

Video of stills shown in Fig. 2B. An S2 cell co-transfected with Draper-GFP, Shark-TagBFP, and Crk-mCherry was allowed to settle on 10% PS supported lipid bilayers and imaged by TIRF microscopy. Cell was imaged every 15 sec. Draper-GFP, Shark-TagBFP, and Crk-mCherry co-localize at mobile microclusters that flow towards the central synapse. Time in min:sec. Scale bar, 5 µm.

**Video 4: Visualization of Draper and F-actin on 10% PS bilayers**

An S2 cell co-transfected with Draper-GFP and Lifeact-TagBFP (Fig. 2D) was allowed to settle on 10% PS supported lipid bilayers and imaged by TIRF microscopy. Cell was imaged every 55 sec. Note that Draper-GFP and Lifeact-TagBFP appear at the plasma membrane together but as clusters mature the two reporters segregate from one another. A prominent F-actin “hole” is visible later in the movie, where the central synapse of Draper-GFP signal appears to exclude F-actin as visualized by Lifeact-TagBFP. Time in sec. Scale bar, 5 µm.

**Video 5: Visualization of FKBP-Bead engulfment by a chimeric FRB-Draper receptor**

An S2 cell, co-transfected with FRB-EXT-Draper-INT-mCherry and Lifeact-eGFP, was incubated with 2% DGS-Ni-NTA/0.5% atto390-PE 6.46 µm silica beads incubated with 10 nM His_10_-FKBP, washed, and resuspended prior to imaging. FKBP beads were incubated with co-transfected S2 cells in the presence of 1 µM rapamycin, allowed to settle in the imaging chamber for 5 min, and visualized at 5 min intervals by spinning disk confocal microscopy. Time in min. Scale bar, 5 µm.

**Video 6: Visualization of FRB-EXT-Draper-INT and Lifeact an FKBP supported lipid bilayer**

An S2 cell, co-transfected with FRB-EXT-Draper-INT-mCherry and Lifeact-eGFP, was incubated with 2% DGS-Ni-NTA/0.5% atto390-PE 6.46 µm silica beads incubated with 1 nM His_10_-FKBP, washed, and resuspended prior to imaging. FKBP beads were incubated with co-transfected S2 cells in the presence of 1 µM rapamycin, allowed to settle in the imaging chamber and imaged at 65 second intervals. Time in sec. Scale bar, 5 µm.

## Supplementary Figure Legends

**Figure S1 (related to Figure 1):**
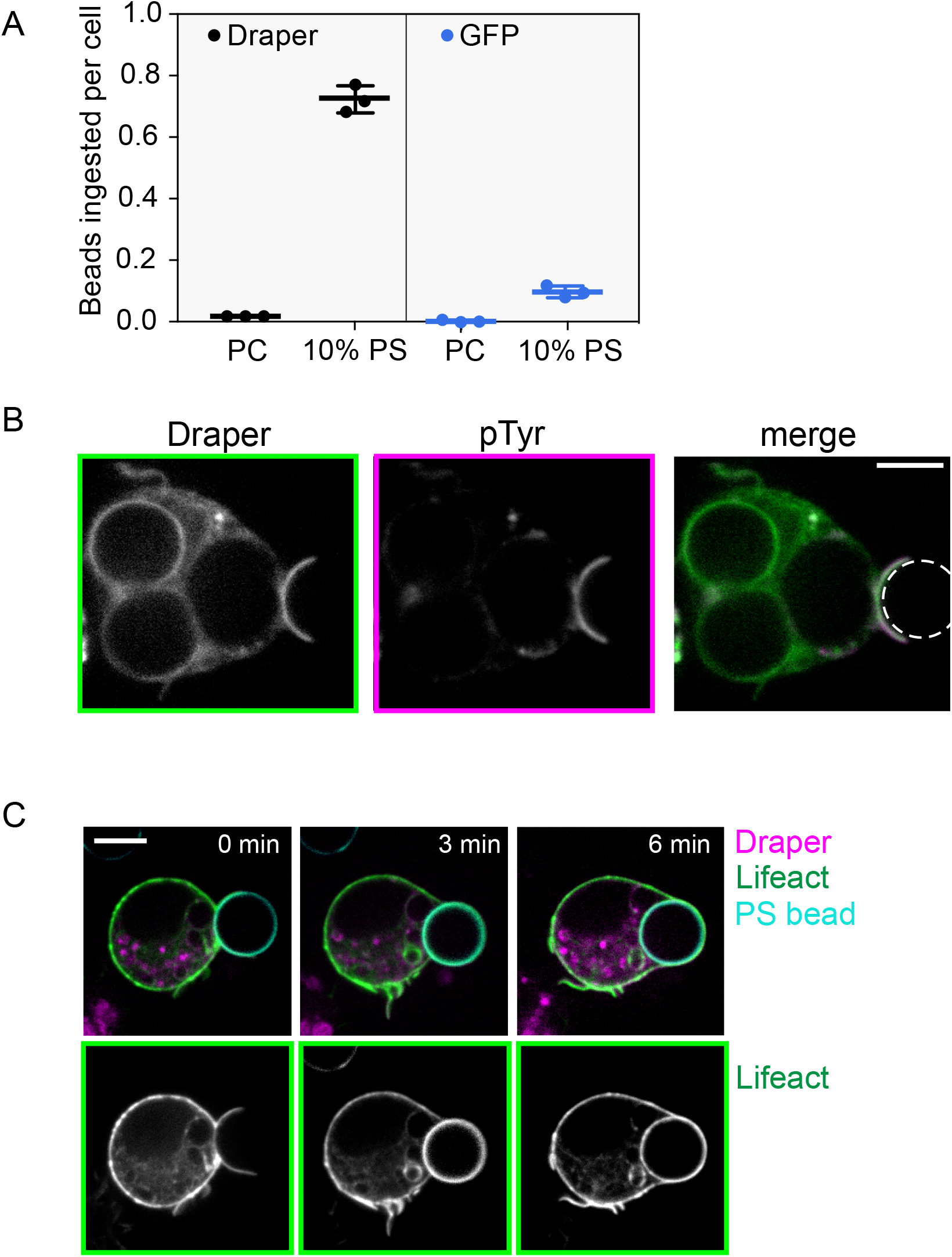
Efficient engulfment of PS beads requires Draper and initiates local signaling. **A.** Draper overexpression increases PS-bead phagocytic proficiency ∼7 fold. S2 cells were transfected with either Draper-GFP or a GFP control vector and incubated with PC or 10% PS beads for 30 min at 27° C. The mean number of beads fully ingested per cell for three independent biological replicates are shown (mean ± SD). **B.** Draper-GFP is tyrosine phosphorylated at the interface with a 10% PS bead. Draper-GFP+ S2 cells were incubated with 10% PS beads. After 30 min incubation cells were fixed, permeabilized and stained with phosphotyrosine (pTyr) antibody. pTyr staining is enriched at the synapse between bead and cell. During fixation and permeabilization, the supported lipid bilayer on the bead is stripped off; bead position is indicated by dashed white line. **C.** F-actin projections around a PS bead. Upon completion of the phagocytic cup, beads are internalized. Draper-GFP/Lifeact-BFP S2 cells were incubated with 10% PS/0.1% rhodamine-PE beads and imaged every 3 min by spinning disk confocal microscopy. A middle section from a z-stack Video is shown. Stills are taken from Video 1. Scale bar indicates 5 µm.

**Figure S2 (related to Figure 2):**
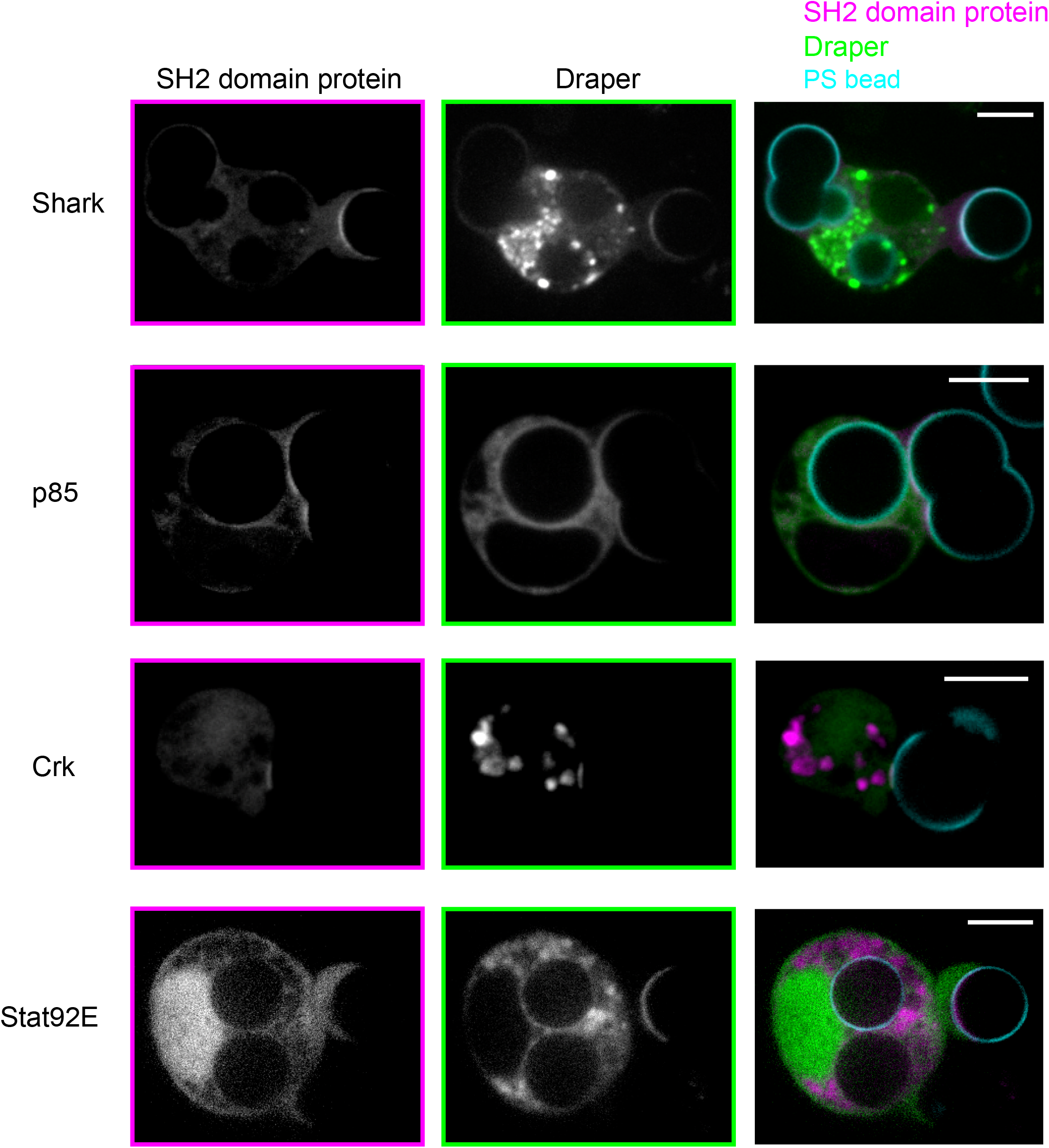
p85 and Crk are recruited to the synapse with a 10% PS bead. Draper-mCherry’s ability to bring the indicated SH2-GFP proteins to the plasma membrane at the synapse with a 10% PS bead. Shark, p85 (the PI3K activating subunit), and Crk were recruited to Draper-mCherry at the synapse, while Stat92E was not recruited. Scale bars indicate 5 µm.

**Figure S3 (related to Figure 2):**
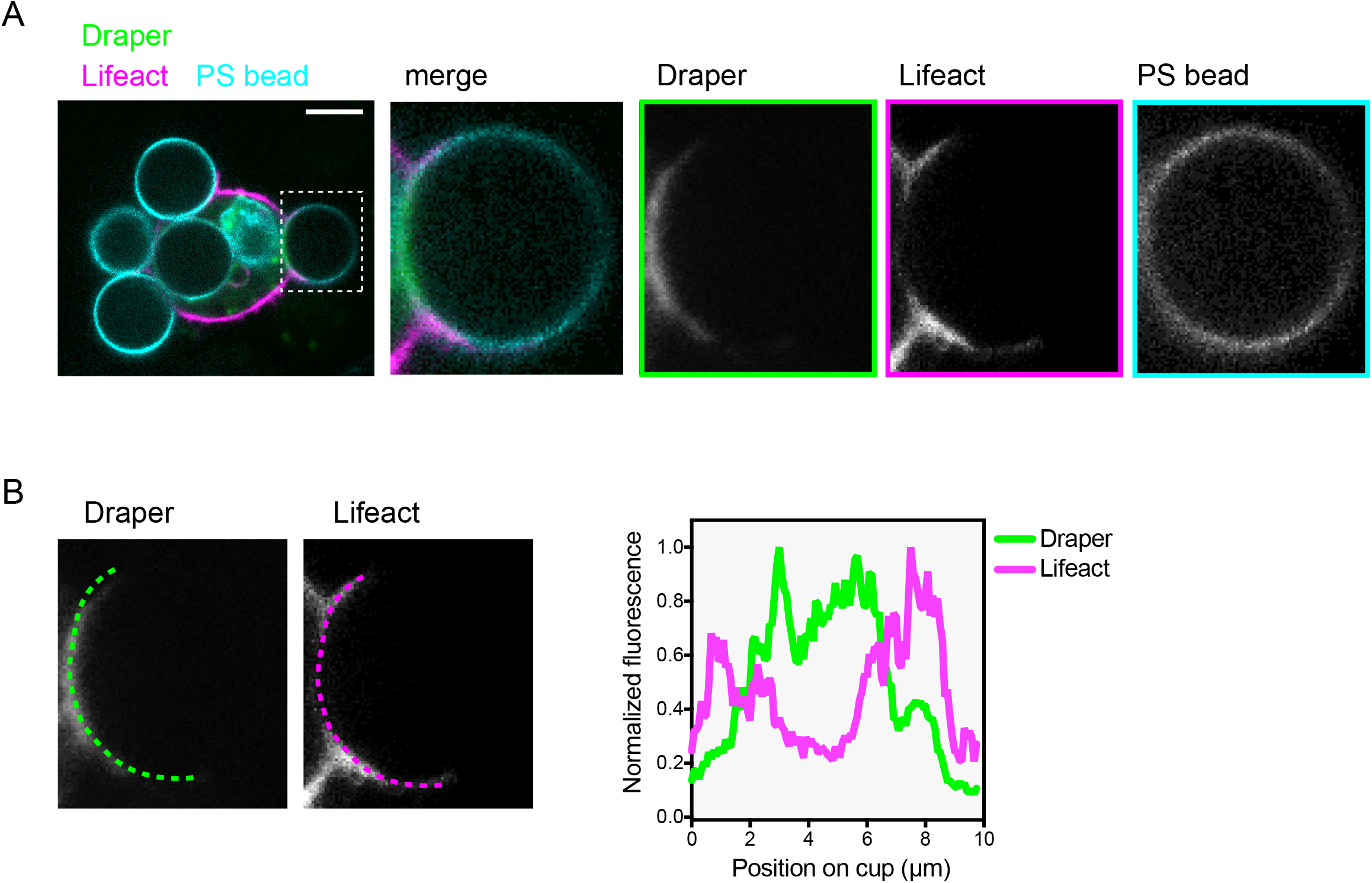
F-actin is depleted at the synapse between PS bead and Draper-expressing S2 cell. F-actin, as measured using a Lifeact-eGFP reporter, is reduced at sites of Draper-mCherry accumulation at the interface between bead and target. Localization of Draper-mCherry and Lifeact-eGFP was assessed after 15 min cell settling by spinning disk confocal microscopy. A 2-pixel linescan of the phagocytic cup was used to measure Lifeact-GFP and Draper-mCherry signal. Green and magenta dashed lines indicate position of linescan along the cup. Fluorescence above background was normalized to the highest value on the linescan. Scale bar indicates 5 µm

**Figure S4 (related to Figure 4):**
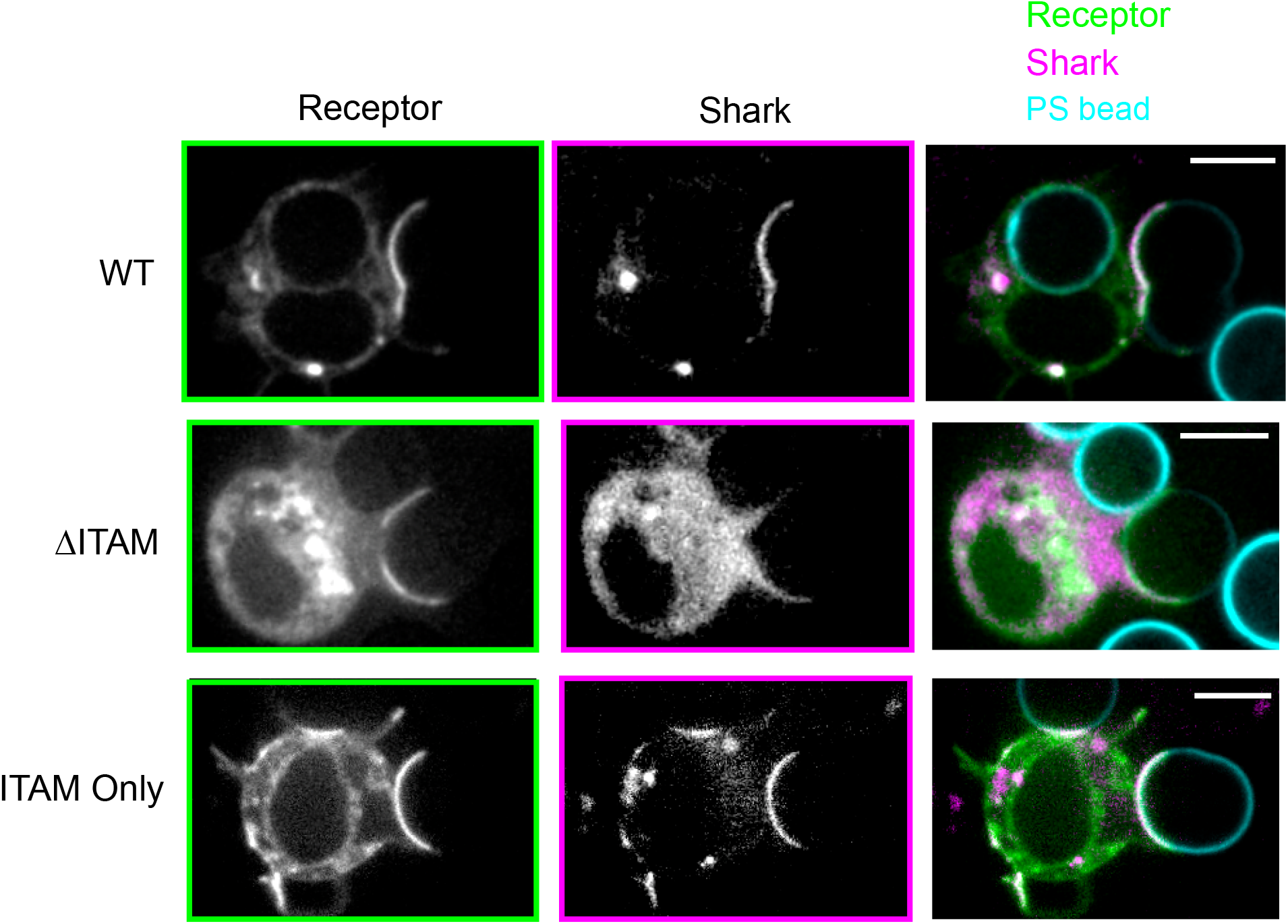
Draper’s ITAM is necessary and sufficient to recruit Shark in cells. S2 cells co-transfected with Draper, Draper-ΔITAM, or Draper-ITAM Only-GFP and Shark-TagBFP were incubated with 10% PS beads labeled with 0.5% atto647-PE. Localization of receptor and Shark-TagBFP was assessed after 15 min cell settling by spinning disk confocal microscopy. Images were acquired using the same microscope settings and scaled equally for comparison. Scale bars indicate 5 µm.

**Figure S5 (related to Figure 4):**
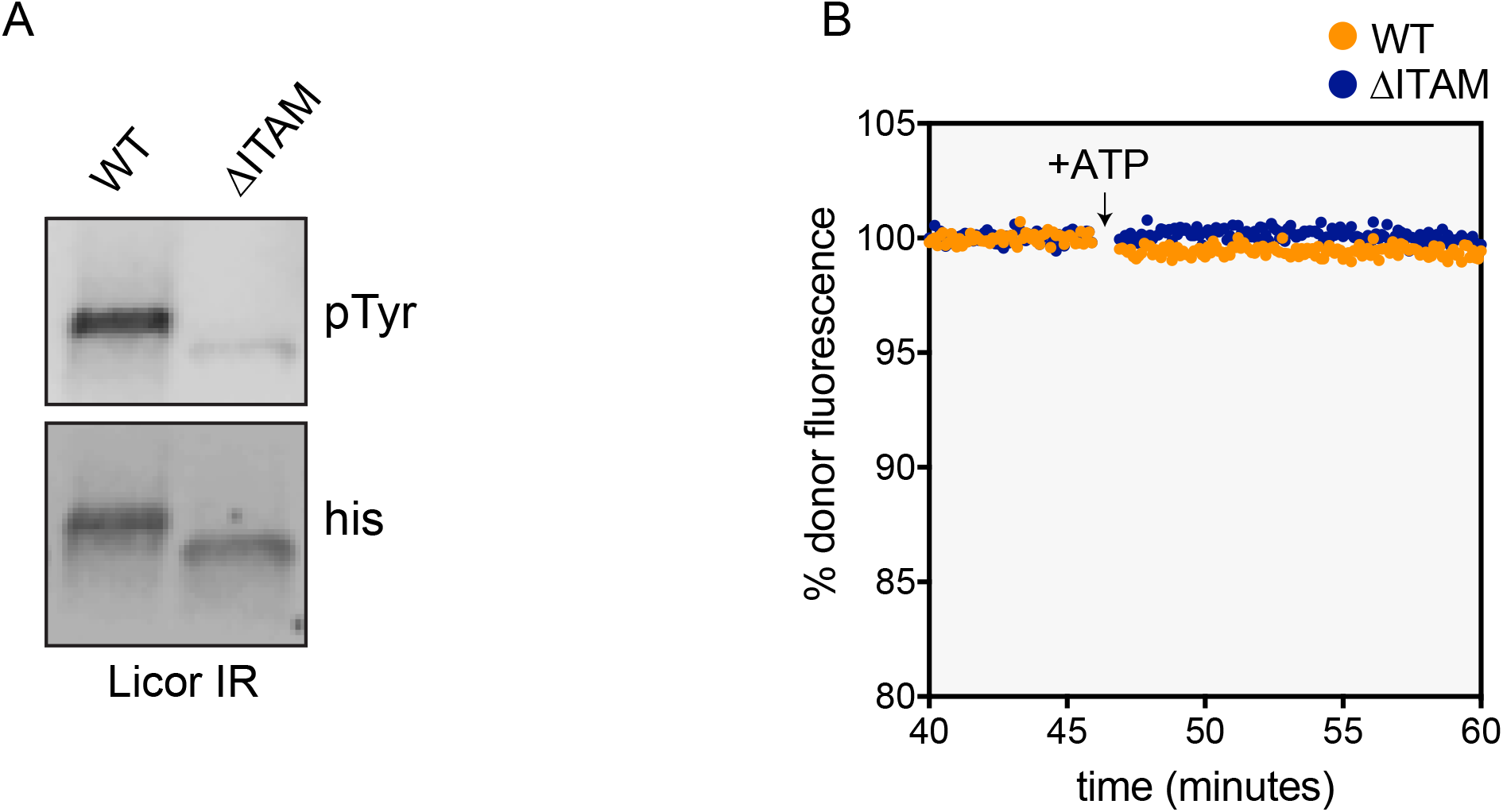
Biochemical reconstitution of Draper phosphorylation reveals a general triggering mechanism and specific effector recruitment. **A.** To perform on-liposome phosphorylation reactions to assess Draper activation by Lck, 1 µM 10his-Draper cytoplasmic domain or 10his-DraperY934F/Y949F (ΔITAM) receptor tail were ligated to liposomes doped with DGS-Ni-NTA and incubated with 1 nM his10-Lck-Y505F and 1 mM ATP for 30 min. Reactions were quenched with SDS-PAGE buffer containing DTT and 2-Me and boiled for 10 min at 95°C. Samples were run on a 4-20% gradient gel and pan phosphotyrosine (800 nm anti-mouse secondary) was assessed by western blotting and Licor imaging. **B.** The Draper cytosolic domain does not directly bind p85^NtSH2^. A BG505 labeled reporter protein for p85^NtSH2^ was prepared as Fig. 4B. His_10_-Draper cytosolic domain or His_10_-DraperY934F/Y949F (ΔITAM) receptor tail were ligated to 0.5% rhodamine-PE liposomes and ATP dependent quenching was assessed. A lack of quenching indicates undetectable direct binding between the p85 reporter construct and both Draper and Draper Y934F/Y949F (ΔITAM).

## Materials and methods

**Note:** *Catalog information for reagents and tools used in this study can be found in a table at the end of this section.*

### Drosophila S2 Cell Culture

Drosophila S2U cells were cultured in Schneider’s media (Gibco) supplemented with 10% Fetal Bovine Serum (FBS, Atlanta) and 1x antibiotic-antimycotic (Gibco). S2 cells were grown in non-vented T25 or T75 flasks (Corning) and split every 3-6 days from high density cultures (∼50-80%). S2U cells were not split to below ∼25% confluency as S2U cells are unhappy at low density. Healthy S2U cells adhere gently to the surface of the culture dish rather than floating, but can be split by gently resuspending the adhered cells using a tissue culture pipet and a 10 ml sterile tissue culture pipet. Enzymatic dissociation using trypsin or by scraping is unnecessary for S2U cells. Frozen stocks were prepared regularly. S2U freeze media is: 45% fresh complete media, 45% sterile filtered conditioned media prepared after spinning the media for 5 min at 1000 x g, and 10% DMSO (Sigma). A confluent T75 dish of S2U cells was pelleted for 5 min at 1000 x g, resuspended in 10 ml of freeze media (described above) and split as 1 ml aliquots in externally-threaded cryotubes (Corning), stored at −80°C in a cell freezing apparatus containing isopropyl alcohol, and transferred to long term storage in liquid nitrogen, vapor phase. Cells were regularly tested for mycoplasma using the MycoAlert kit (Lonza).

### Transfection of Drosophila S2U Cells

Cells were split to ∼60% confluency into 6 well tissue culture dishes 4 to 12 hr prior to transfection. All transfection mixes were prepared in 5 ml low-retention polystyrene round-bottom tubes. To prepare transfection mixes a maximum of 2.5 µg plasmid DNA was added to 300 µl serum-free Schneider’s media and gently mixed. 5 µl of TransIT-Insect (Mirus) was added to diluted DNA and gently mixed. At this point the solution appears cloudy. Transfection mixes were incubated at room temperature for 15 min and added drop-wise gently to the 6 well plate using a p1000 pipet. After drop-wise addition of transfection mixes, the plate was gently shaken laterally to ensure even distribution of TransIT-DNA particles. Transfection efficiencies of at least 50% and up to 90% were regularly obtained using this method, an improvement compared to other transient transfection methods employed for Drosophila S2U cells.

All constructs used in this study express proteins of interest in S2U cells were under control of a pMT copper-inducible promoter. To induce expression in transfected cells, CuSO_4_ was added from a 50 mM sterile stock in water to a final concentration of 250 µM (1 to 200 dilution from stock) 36 to 48 hr after transfection. The timing of induction is important – at this later time point, we find expression levels to be lower and more uniform. At shorter time points (e.g. 12 to 24 hr after transfection), transfection efficiency was lower and expression levels in transfected cells higher, as assessed by live cell imaging of fluorescent proteins under control of the pMT promoter. Detailed information about all constructs used in this study is available in Supplemental Excel Table 1. For the experiments involving Draper-GFP or –mCherry overexpression, the long activating isoform Draper-I (1031 amino acids, NP_728660.2) was used.

### Preparation of APC Annexin V Labeled Apoptotic Cell Corpses

To prepare labeled apoptotic cell corpses for use in internalization assays, 1 µM final concentration of Actinomycin D (Sigma; 1 mg/ml in DMSO, 833 µM stock) was added directly to 80% confluent cultures of Drosophila S2U cells in 6 well plates. Cells were incubated for 16 hr in Actinomycin D prior to harvesting by centrifugation for 5 min at 1000 x g in a 15 ml falcon tube. Actinomycin D was washed out by rinsing corpses twice in complete Schenider’s media in the same falcon tube – sequential washes, each with 10 ml complete media were performed. Corpses from 1 well of a 6 well dish were resuspended in 300 µl of complete media after washes and transferred to a 1.7 ml Eppendorf tube for labeling. To label externalized phosphatidylserine (PS), APC Annexin V (BioLegend) was added directly to the washed corpses to a final concentration of 50 ng/ul (from 5 µg/ml stock). The tube was wrapped in foil prior to labeling on a rotator at room temperature for 30 min. After labeling, cells were washed 3 times by sequential harvesting and resuspension in complete media. After washing labeled corpses were resuspended in 300 µl complete media.

### Assay for Internalization of Apoptotic Cell Corpses

To assess internalization of apoptotic cell corpses, washed Draper-GFP or mCherry-control transfected S2U cells were incubated with fresh complete media containing 10% FBS and labeled cell corpses prepared as above. Transfected cells were washed with complete S2U media, and culture media was replaced with 1 ml of fresh. Cells in fresh media were gently resuspended by pipetting up and down. 10 µl of the washed labeled cell corpses prepared as above were mixed in an Eppendorf tube with 300 µl of washed transfected cells. Eppendorf tube was flipped up and down 4 times and complete mixture was plated in 1 well of a 96-well glass bottomed MatriPlate imaging dish (Brooks). After 45 min incubation at room temperature in the dark, High Content Screening imaging and quantification were performed as described below.

### Preparation of 10% PS-coated Silica Beads

10 molar % phosphatidylserine (PS) beads were used for all experiments other than the PS titration in Figure 1G. To prepare 10 molar % beads, chloroform suspended lipids sufficient for 1 ml of a 10 mM solution were mixed at the following proportions: 89.5% POPC, 10% DOPS, 0.5% PEG5000-PE, 0.5% atto390-DOPE (label for visualization by microscopy). Lipids were transferred to a chloroform-washed glass vial using gas-tight Hamilton syringes and dried to a film under argon gas in a warm (∼45° C) water bath. Dried lipids were stored under foil (if labeled lipids were used) and desiccated overnight at room temperature in a benchtop desiccator filled with Drierite. Dried lipids were suspended in 1 ml tissue culture grade PBS by vortexing for 1 min under parafilm and gentle pipetting. Suspended lipids were transferred to a 1.7 ml Eppendorf tube and stored under argon gas. The PBS-lipid mix was freeze/thawed 5 times in liquid nitrogen/warm water and subjected to 2 x 5 min sonication in a Bioruptor Pico bath sonicator (Diagenode). Sonicated lipids were spin at 35000 rpm for 35 min at 4° C in a TLA 120.1 rotor in a benchtop Beckman ultracentrifuge. Only a small pellet should be visible after spinning sonicated lipids. Small Unilamellar Vesicles (SUVs) prepared using this method were used immediately or stored under argon gas, flash frozen, and stored at −80° C. Lipid mixes are best used within two weeks of preparation. Undiluted 10 mM lipid mixes were used as described below to build bilayers on silica beads. For the PS titration in Figure 1G differences in PS concentration were made up by balanced changes to the molar % of PC such that final lipid concentration in 1 ml PBS matched the 10 mM concentration described above.

50 µl 6.46 µm silica beads (Bangs, Catalog #SS06N) were added to 300 µl Milli-Q water in 1.7 ml Eppendorf tubes for washing. Beads in suspension were pelleted 3 times in 300 µl Milli-Q water at 300 rcf and decanted. After the third wash, beads were resuspended in ∼30 µl PBS until the pellet was barely covered. 300 µl of 2 mM final concentration of the desired lipid mix (diluted from 10 mM stock in PBS) was vortexed briefly, wrapped in foil, and rotated at room temperature for 45 min. Beads were pelleted and washed 3 times by sequential pelleting and resuspension in 300 µl fresh PBS. Washed beads were resuspended in 300 µl fresh PBS for bead assay, described below.

### Assay for Internalization of PS-coated Silica Beads

To assess internalization of PS-coated silica beads, 7 µl of bead suspension was mixed with 300 µl washed transfected S2U cells resuspended in fresh complete media containing 10% FBS, flipped gently to mix 4 times, and the complete mixture plated in 1 well of a 96-well glass bottomed MatriPlate imaging dish (Brooks). Cells were allowed to settle and imaged as described below. For endpoint assays, quantification was performed on images taken 30 or 45 min after plating. For kinetic experiments, imaging was started 5 min after plating. Engulfment experiments were performed using S2U cells co-transfected with GFP or a GFP-fusion receptor protein, mCherry-CAAX (full-length mCherry-LEKMSKDGKKKKKKSKTKCVIM) for visualizing plasma membranes during quantification (described below), and 10% PS atto390-PE labeled silica beads.

### Assay for Internalization of His_10_-FKBP Silica Beads

Engulfment experiments using His_10_-FKBP ligated beads were performed as above with the exception that FRB-EXT-Draper-INT-GFP expressing S2 cells were co-transfected with mCherry-CAAX to serve as a specific receptor for the FKBP ligand. Silica beads above were prepared as above with the exception that instead of the 10% PS lipid mix, the following lipid mix was used: 97% POPC, 2% DGS-NTA-Nickel, 0.5% PEG5000-PE, 0.5% atto390-DOPE. Beads were blocked for 15 min in 300 µl PBS + 0.1% w/v BSA. His_10_-FKBP was diluted to 10 nM final concentration in PBS + 0.1% w/v BSA and 300 µl of this dilution was added to each 50 µl bead pellet, as described above. Proteins were coupled to beads for 40 min under foil on a rotator at room temperature, washed 3 x 5 min in PBS + 0.1% w/v BSA and resuspended in PBS + 0.1% w/v BSA prior to the engulfment assay. 300 µl washed transfected cells were mixed with 7 µl His_10_-FKBP ligated beads either in the presence of 1 µM rapamycin or a matching volume of DMSO (vehicle), flipped 4 times to mix, and ingestion was quantified as described below.

### Staining for Phosphotyrosine (pTyr) and the Synapse between S2 Cell and Bead

To fix and stain bead-cell synapses in chambers described above for phosphotyrosine (pTyr) localization, half the media (∼150 µl) from imaging chambers was gently removed and replaced with 150 µl 6.4% paraformaldehyde. Cells were fixed under foil for 15 min, washed 2 x 3 min in PBS, permeabilized with 0.5% v/v Triton X-100 in PBS for 10 min, and set overnight to block in PBS + 0.1% v/v TritonX-100 + 0.2% w/v BSA at 4° C wrapped in parafilm in the dark. Cells were stained in primary antibody (pY-20 mouse anti-phosphotyrosine (Santa Cruz), 2 μg/ml final concentration, 1:100 dilution) for 1 hr at room temperature in the dark, washed 3 x 5 min room temperature in PBS + 0.1% v/v Triton X-100 + 0.2% w/v BSA, and incubated in secondary antibody (Thermo, anti-mouse IgG H+L Alexa Fluor 647 conjugated) for 1 hr at room temperature, washed 3 x 5 minutes in PBS + 0.1% v/v Triton X-100 + 0.2% w/v BSA. Washed fixed stained cells were gently covered in 200 µl PBS prior to imaging. Cells were imaged immediately or stored for up to 2 days at 4° C in the dark, wrapped in parafilm before imaging my spinning disk confocal microscopy (see below).

### Building PS-containing Supported Lipid Bilayers (SLBs)

The night before the experiment, a 96 well MatriCal imaging plate was submerged in 2% v/v Hellmanex III, microwaved for 2.5 min on full power, and set on a stir plate under saran wrap for at least 12 hr. Hellmanex was washed away through 20 sequential washes using Milli-Q water and the washed plate dried under Nitrogen gas. The dried plate was covered in thermal paper using a roller. The desired number of wells were exposed using a razor and washed 3 x 5 min 5N NaOH. After NaOH washes, wells were cleaned 10 times with MilliQ water.

To prepare 10% PS bilayers, 250 µl 2 mM lipid mix in PBS described above gently pipetted on the glass surface. The lipid mix was pipetted up and down and bilayer formation was allowed to proceed for 30 min at room temperature. After bilayers formed, washes were performed using PBS 3 x 5 min. For each wash, PBS was gently pipetted up and down over the surface of the bilayer to remove SUVs not part of the SLB. Bilayers were washed into serum free Schneider’s media prior to adding cells. Imaging on bilayers should be complete within 45 min for best results, as bilayer integrity cannot be assumed after this time.

After washing the SLB, 150 µl of washed cells were added to the surface, allowed to settle, and imaged by Total Internal Reflection Fluorescence-Microscopy as described below. A lower density of cells can be used, as cells will spread on bilayers and imaging overlapping cells is not ideal.

### Building His_10_-FKBP Bearing Supported Lipid Bilayers (SLBs)

Bilayers were prepared as above with the exception that the following lipid mix was used (note the use of DGS-NTA-Nickel lipid to ligate His_10_ proteins to the bilayer): 97% POPC, 2% DGS-NTA-Nickel, 0.5% PEG5000-PE, 0.5% atto390-DOPE or 0.5% atto647-DOPE for labeling, depending on the fluorescent proteins present on fusion proteins assessed. Washed bilayers were blocked in PBS + 0.1% w/v BSA for 15 min prior to protein coupling. To couple His_10_-FKBP to 2% DGS-NTA-doped bilayers, His_10_-FKBP was diluted to 10 nM final concentration in PBS + 0.1% w/v BSA and added to SLBs to couple for 40 min at room temperature. Uncoupled protein was washed 3 x 5 min under foil at room temperature prior to adding cells as above with the exception that when FKBP ligand was used with receptors fused to FRB-extracellular domains, 1 µM rapamycin was added to form the 100 fM FRB-rapamycin-FKBP ternary complex.

### Liposome FRET Assay for Effector Recruitment to Phosphorylated Receptor Tails

This assay was based upon one used to reconstitute mammalian T cell signaling (Hui and Vale, 2014). To present receptor tails in physiological geometry for phosphorylation by Src-family kinases, 1 µM His_10_ receptor tails (either His_10_-Draper-INT or His_10_-Draper-ΔITAM-INT) were ligated to liposomes comprised of the following molar percent: 74.5% POPC, 10% DGS-NTA, 0.5% Rhodamine-PE, 15% DOPS. Liposomes were prepared using an extruder with 200 nm filters (Avanti). Receptor tails and 1 nM His_10_-Lck-Y505F or 86 nM Src42a (soluble) were equilibrated with receptor tails in the presence of BG505-kinase-tSH2 reporter proteins were incubated. After proteins were allowed to bind for 40 min at room temperature in the dark. At this point, Mg-ATP was added to a 1 mM final concentration and BG505 signal was assessed at indicated time points. Quenching of BG505 signal, dependent upon addition of ATP, indicates recruitment of reporters to phosphorylated receptor tails. Quenching of the BG505 dye was assessed using a Synergy H4 plate reader (BioTek).

### Determining Sites of Tyrosine Phosphorylation by 2D Mass Spectrometry

To determine sites of Tyr phosphorylation on Draper (Figure 6), 1 µM His_10_-Draper-INT was ligated to unlabeled liposomes: 74.5% POPC, 10% DGS-NTA, 15% DOPS and phosphorylated by 1 nM His_10_-Lck-Y505F after 40 min of protein coupling to liposomes. 100 µl Reactions were quenched with 100 µl of 8 M Urea and 20 mM EDTA at the indicated time points (final concentrations of 4 M urea and 10 mM EDTA) and shock frozen in liquid nitrogen. To perform trypsinization, reactions were thawed by boiling, supplemented with 5 mM final concentration of TCEP, and incubated for 20 min at room temperature. After denaturation samples 10 mM final concentration of iodoacetamide (IAA) as added. Samples were then incubated in the dark for 15 min. Urea concentration was reduced two-fold through addition of 200 µl PBS. 1 mM final concentration of CaCl_2_ and 1 µg/ml Trypsin Gold (Promega) was added before samples were incubated overnight in the dark at 37° C. Trypsinzation was quenched through addition of 50 µl mass spec grade formic acid. Samples were then subjected to 2D mass spectrometry to remove contaminants and peptides containing the characteristic shift for phosphorylation (+79.9663 Da) quantified and expressed as a percentage of all peptides covering the indicated residue.

### Immunoblot Analysis

To determine degree of tyrosine phosphorylation on His_10_-Draper intracellular domains, on liposome phosphorylation assays were performed as above and quenched using SDS-PAGE buffer + DTT and incubated for 5 min at 95°C. Reactions were run on 4-20% gradient SDS-PAGE gels and transferred onto nitrocellulose membranes using the iBlot system (Invitrogen). Membranes were blocked for 1 hr at room temperature in PBS + 0.1% w/v bovine serum albumin (BSA, Sigma) and incubated overnight at 4°C with anti-His probe (rabbit) and anti-phosphotyrosine (mouse) antibodies (Key Resources Table). Primary antibodies were incubated with membranes simultaneously. Both were used at 1:1000 final dilution PBS + 0.1% w/v BSA. Membranes were washed 3 x 5 minutes at room temperature in PBS + 0.1% w/v BSA and incubated with Licor secondary antibodies at 1:10,000 dilution (700 nm anti-rabbit, 800 nm anti-mouse, Key Resources Table) for 1 hr at room temperature. Membranes were washed 3 x 5 minutes at room temperature in PBS + 0.1% w/v BSA and imaged using a Licor Odyssey gel imager. Quantification of anti-phosphotyrosine and anti-his intensity was performed using the Gel Analysis feature of Fiji.

### Protein Expression, Purification, and Labeling

His_10_-Draper and His_10_-Draper-ΔITAM were transformed into BL-21 DE3 RIPL cells (Agilent). Cells were grown in Terrific Broth (TB), 12 g tryptone, 24 g yeast extract, 4 ml glycerol up to 1L in MilliQ water + 10% v/v sterile salt solution in MilliQ water: 0.17M KH_2_PO_4_, 0.72 M K_2_HPO_4_, until cells reached an OD_280_ of ∼0.6-0.8, chilled in a 4°C room, and induced overnight by addition of 0.5 mM final concentration of IPTG. Protein expression was performed in shaker set to 18°C. Bacterial pellets were resuspended in lysis buffer (50 mM HEPES, pH 7.5, 300 mM NaCl, 1 tablet Complete EDTA-free protease inhibitor tablet crushed per 100 ml buffer). Lysates were prepared by probe sonication and bound to Ni-NTA agarose (Qiagen). Beads were washed extensively with lysis buffer and proteins eluted in 500 mM imidazole in lysis buffer. Imidazole elutions were subjected to gel filtration chromatography using a Superdex 200 10/30 column (GE Healthcare) in gel filtration buffer: 50 mM HEPES, pH 7.5, 150 mM NaCl, 10% glycerol, and 1 mM TCEP. Monomer fractions were pooled, shock frozen in liquid nitrogen, and stored in small aliquots at −80°C. Soluble Src42a was purified from SF9 cells as a GST-(PP)-SNAP-Src42a fusion where PP indicates a cut site for Precission Protease. Precission Protease cleavage was performed overnight and post-cleavage supernatants were gel filtered as described below. His_10_-Lck Y505F was purified from SF9 cells over Nickel agarose (Qiagen) and eluted in 500 mM imidazole in lysis buffer. Both Src family kinases were subjected to gel filtration chromatography using a Superdex 200 10/30 column (GE Healthcare) in gel filtration buffer: 50 mM HEPES, pH 7.5, 150 mM NaCl, 10% glycerol, and 1 mM TCEP. Monomer fractions were pooled, shock frozen in liquid nitrogen, and stored in small aliquots at −80°C. The tandem SH2 domain reporter for Shark kinase was cloned as a GST-(PP)-SNAP-Shark-tSH2 fusion where PP indicates a cut site for Precission Protease. The soluble SNAP-Shark-tSH2 reporter was generated by on-resin cleavage with Precission Protease during purification. GST-Precission Protease remained on the beads. Cleavage products were subjected to gel filtration chromatography using a Superdex 200 10/30 column (GE Healthcare) in gel filtration buffer: 50 mM HEPES, pH 7.5, 150 mM NaCl, 10% glycerol, and 1 mM TCEP. To prepare BG505-labeled SNAP-Shark-tSH2 for the liposome FRET assay, 10 µM cleaved, gel filtered monomer was incubated at a 1:2 molar ratio with SNAP-Cell 505 Star (NEB) overnight in the dark at 4°C and run over a PD Minitrap G-25 column (GE Healthcare) to eliminate excess dye.

### Spinning Disk Confocal Microscopy

All internalization and bead co-localization imaging performed for this study was done of a spinning disk confocal microscope (Nikon Ti-Eclipse inverted microscope with a Yokogawa spinning disk). For bead internalization kinetic assays images were acquired using a 40x/0.95 N/A air objective. Live image acquisition for internalization of corpses and beads was performed using the High Content Screening (HCS) Site Generator plugin in µManager. All other images were acquired using a 100x 1.49 N/A oil immersion objective. Images were captured using an Andor iXon EM-CCD camera. The open source µManager software package was used to run the microscope and capture the images (Edelstein et al., 2010). All raw microscopy images were acquired as 16-bit TIFF stacks.

### Total Internal Reflection Fluorescence-Microscopy (TIRF-M)

All SLB imaging was performed using a TIRF microscope equipped with a motorized TIRF arm and a Hamamatsu Flash 4 camera. For all TIRF-M imaging images were captured at 2 x 2 binning. The open source µManager software package was used to run the microscope and capture the images (Edelstein et al., 2010).

### Imaging processing and analysis

All image quantification was done on raw, unprocessed images. All images in figures were opened in ImageJ. A middle z-slice was extracted and the channels split. The image intensities were scaled to enhance contrast in Photoshop (Adobe) using appropriate levels of linear adjustment. Background correction was not performed. Where mutants are compared in the same series, images are acquired using identical microscope and camera settings and scaled to equal intensity values.

### Quantification and Statistical Analysis

For both corpse internalization and bead internalization assays, targets were scored as internalized following complete engulfment of the object, as assessed by visible mCherry-CAAX or receptor-GFP signal visible completely around the target in stills from live imaging. Beads or corpses attached but not internalized were not scored. Only targets with labeled Annexin V or atto390 visible were scored as internalized. At least 150 cells per condition are shown. Mean value and ± one standard deviation are plotted (GraphPad 6.0).

### Quantification and Statistical Analysis – Draper microcluster velocity calculation

Quantification of microcluster velocity was performed in MicroManager (Edelstein et al., 2010). Microcluster velocity was calculated by measuring the distance a single cluster traveled over 150 or 165 seconds of a TIRF-M movie captured at 15 sec intervals. Quantification was performed on the same cell population shown Figure 2A (Draper-GFP, Crk-mCherry, Abl-TagBFP transient transfection settling on 10% PS supported lipid bilayers). A single channel (Draper-GFP) Maximum Intensity Z-projection was created from a substack of 10 or 11 intervals. The freehand line tool was used to measure distance traveled by n=50 microclusters on the Maximum Intensity Z-projection. The reported velocity in the text is the mean for n=50 individual representative microclusters. Mean microcluster velocity observed was 1.92 µm/ min (SD± 0.61), median value 1.77 µm/min.

### Quantification and Statistical Analysis – Quantification of Draper, Shark, and Lifeact intensity

Quantification of normalized fluorescence intensity was performed in MicroManager (Edelstein et al., 2010). Intensity for each indicated channel was collected across a 2-pixel linescan for each indicated time point using the Plot Profile tool in ImageJ. Intensity measurements were performed in the same manner for TIRF-M and spinning disk confocal imaging. Background was measured as the mean value for a 2-pixel linescan at a region adjacent to the cell. Background was subtracted from intensity at each position before normalization. Values were normalized to the maximum intensity across the linescan. Normalized intensity values are plotted using distance along the transect as the x axis. Transects are indicated on inset images for each cluster.

### Summary paragraph for supplemental material

Figures S1-S5 contain supporting microscopy images and information directly in support of the conclusions of this study. Full legends accompany each supplemental figure. Figure S1: supports Figure 1 by demonstrating both a Draper and PS dependence for efficient engulfment as well as initiation of pTyr signaling and projections around a PS bead. Figure S2: supports Figure 2 by showing that the engulfment effectors that localize to signaling microclusters are also recruited to Draper at the synapse with a 10% PS bead. Figure S3: supports Figure 2 by showing that actin depletion behavior is observed in the bead assay. Figure S4: supports Figure 5 by demonstrating that Draper’s ITAM is necessary and sufficient for Shark kinase recruitment in cells. Figure S5: Supports Figure 5 by showing that, like Src42a, the Src family kinase Lck preferentially modifies Draper based on the presence of an ITAM. This figure also supports Figure 2 and Figure 5 by showing that the N-terminal SH2 domain of p85 does not directly bind Draper’s cytoplasmic tail in an on-liposome biochemical assay.

### Summary paragraph for supplemental material

Figures S1-S5 contain supporting data directly in support of the conclusions of this study. Full legends accompany each supplemental figure. Figure S1: supports Figure 1 by demonstrating both a Draper and PS dependence for efficient engulfment as well as initiation of pTyr signaling and projections around a PS bead. Figure S2: supports Figure 2 by showing that the engulfment effectors that localize to signaling microclusters are also recruited to Draper at the synapse with a 10% PS bead. Figure S3: supports Figure 3 by presenting the expression data for transmembrane phosphatases in Drosophila S2 cells. Figure S4: supports Figure 4 by demonstrating that Draper’s ITAM is necessary and sufficient for Shark kinase recruitment in cells. Figure S5: Supports Figure 4 by showing that, like Src42a, the Src family kinase Lck preferentially modifies Draper based on the presence of an ITAM. This figure also supports Figure 2 and Figure 5 by showing that the N-terminal SH2 domain of p85 does not directly bind Draper’s cytoplasmic tail in an on-liposome biochemical assay.

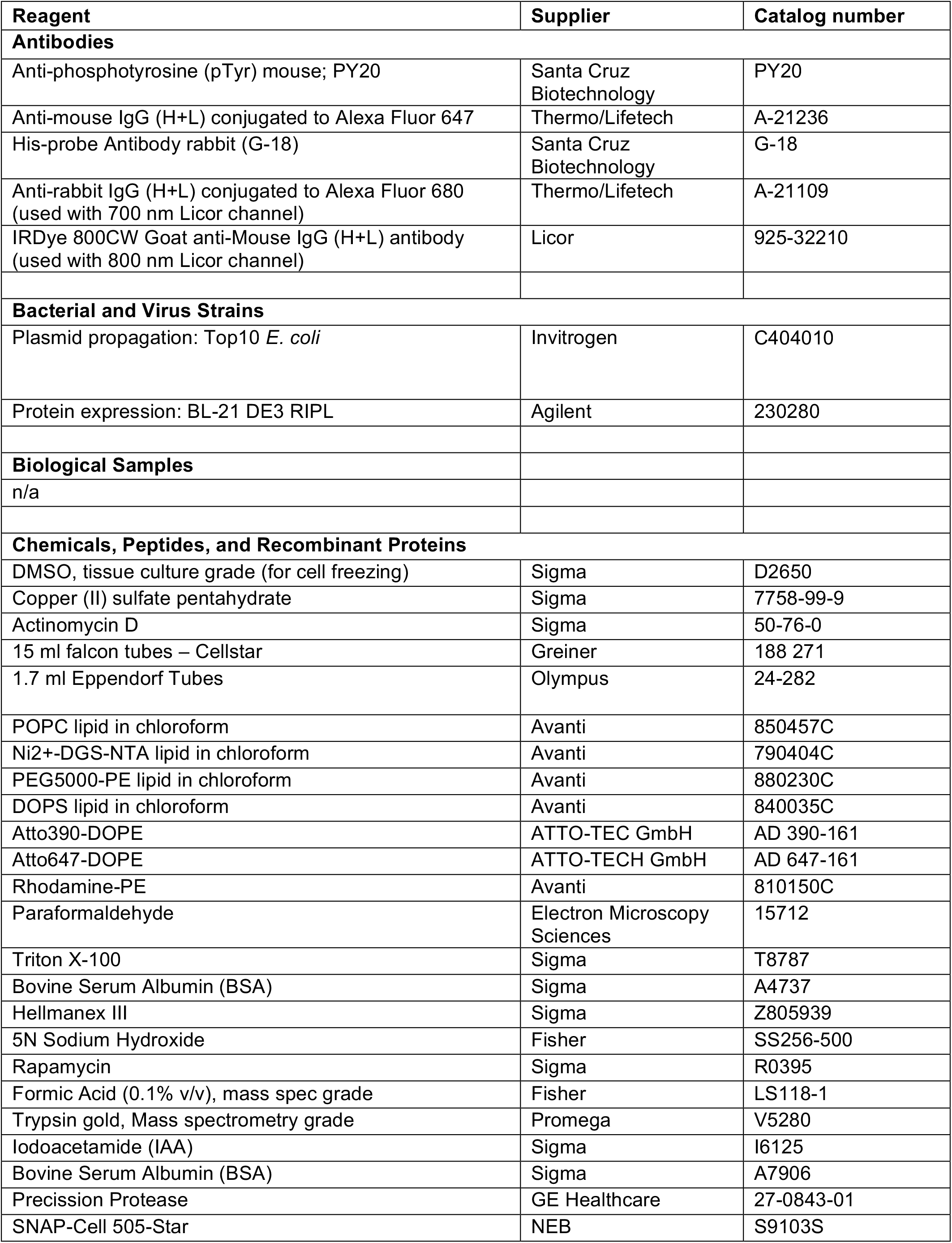
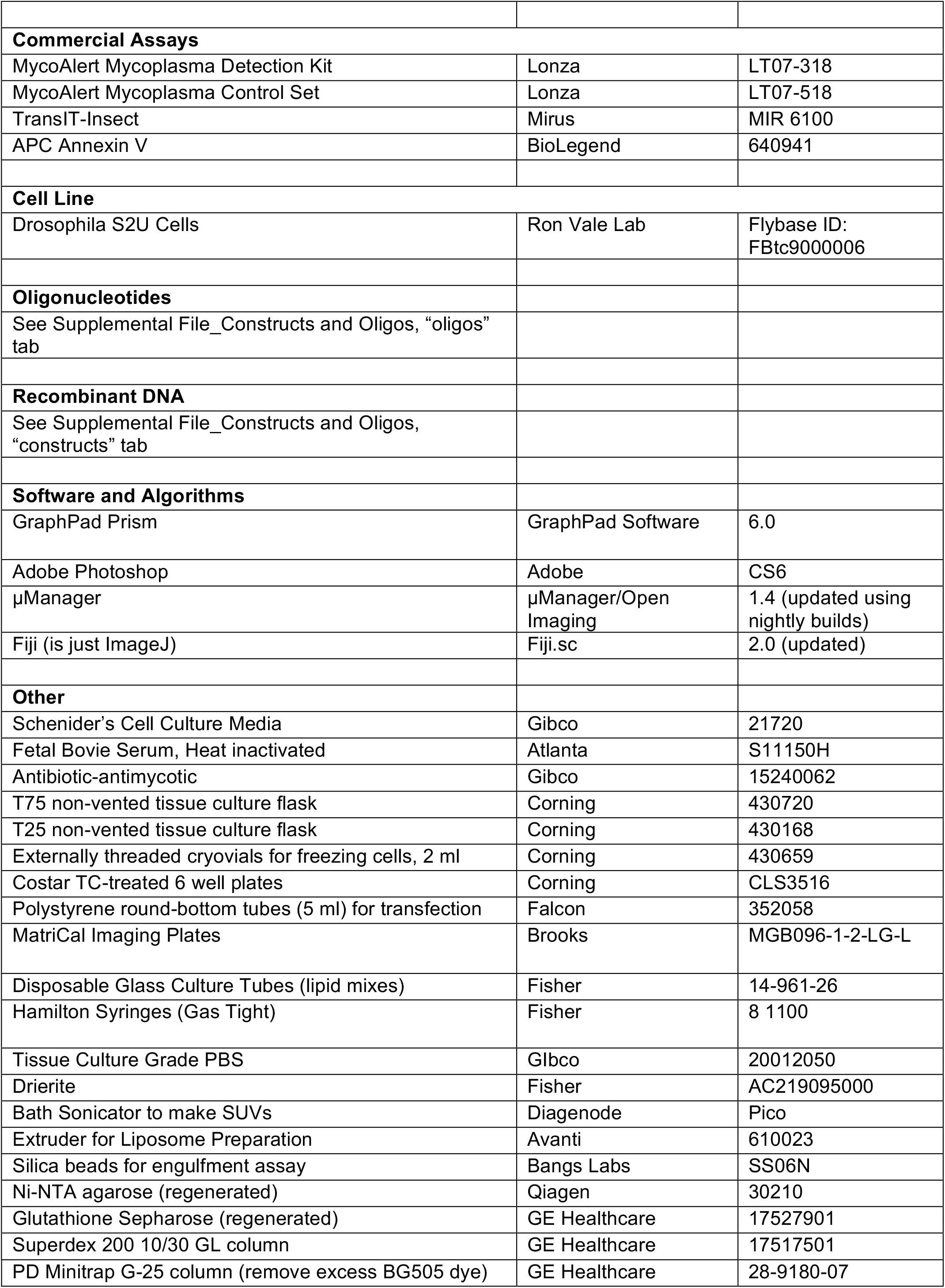
Full supplier and catalog information for materials used in this study.

